# Synthetic genomic reconstitution reveals principles of mammalian *Hox* cluster regulation

**DOI:** 10.1101/2021.07.07.451065

**Authors:** Sudarshan Pinglay, Milica Bulajić, Dylan P. Rahe, Emily Huang, Ran Brosh, Sergei German, John A. Cadley, Lila Rieber, Nicole Easo, Shaun Mahony, Matthew T. Maurano, Liam J. Holt, Esteban O. Mazzoni, Jef D. Boeke

**Affiliations:** Institute for Systems Genetics, NYU Langone Health, New York, NY 10016, USA; Department of Biology, New York University, New York, NY 10003, USA; Center for Eukaryotic Gene Regulation, Department of Biochemistry and Molecular Biology, The Pennsylvania State University, University Park, PA 16802, USA; Department of Pathology, NYU Langone Health, New York, NY 10016, USA; Department of Biochemistry and Molecular Pharmacology, NYU Langone Health, New York, NY 10016, USA; Department of Biomedical Engineering, NYU Tandon School of Engineering, Brooklyn, NY, 11201, USA

## Abstract

Precise *Hox* gene expression is crucial for embryonic patterning. Intra-*Hox* transcription factor binding and distal enhancer elements have emerged as the major regulatory modes controlling *Hox* gene expression. However, quantifying their relative contributions has remained elusive. Here, we introduce ‘synthetic regulatory reconstitution’, a novel conceptual framework for studying gene regulation and apply it to the *HoxA* cluster. We synthesized and delivered variant rat *HoxA* clusters (130-170 kilobases each) to an ectopic location in the mouse genome. We find that a *HoxA* cluster lacking distal enhancers recapitulates correct patterns of chromatin remodeling and transcription in response to patterning signals, while distal enhancers are required for full transcriptional output. Synthetic regulatory reconstitution is a generalizable strategy to decipher the regulatory logic of gene expression in complex genomes.

**One-Sentence Summary:** Reconstitution of gene regulation using large DNA constructs unravels the regulatory logic of a developmental gene locus.

## Introduction

Developmental programs require precise spatial and temporal control of gene expression. To ensure this, developmentally important genes, such as *Hox* genes, are subject to multiple regulatory mechanisms that ensure appropriate expression patterns (*1–3*). *Hox* genes encode evolutionarily conserved transcription factors with crucial roles in cell fate patterning (*4, 5*). Alteration of *Hox* gene expression patterns results in gross developmental defects, even transforming one body part into another (homeotic transformations) (*6*).

*Hox* genes are found within a unique genomic regulatory environment. In mammals, they are organized in four chromosomal clusters (*HoxA, HoxB, HoxC*, and *HoxD*), each harboring a subset of 13 *Hox* paralogs. Each cluster contains *Hox* genes tightly arranged within about 100kb in the same transcriptional orientation and lacks other coding genes (*7*). The spatial and temporal *Hox* gene expression patterns along the anterior-posterior (AP) axis of the developing embryo mirror the organization of the genes within the cluster, a phenomenon known as colinearity (*4, 8*). Thus, the precisely regulated *Hox* clusters have become a paradigm for studying the fundamental link between genome organization and gene expression.

Low gene density regions peppered with distal regulatory elements surround *Hox* clusters. A collection of intricate genetic manipulations has revealed a complex regulatory landscape dispersed within the gene-poor region surrounding the *Hox* clusters (*9–13*). For example, the *HoxA* cluster relies on several retinoic acid (RA) and Wnt responsive distal regulatory elements (enhancers) located in the gene desert between *Hoxa1* and the next gene *Skap2* (∼300kb away) (*14–17*) (Figure 1A).

**Figure 1:**
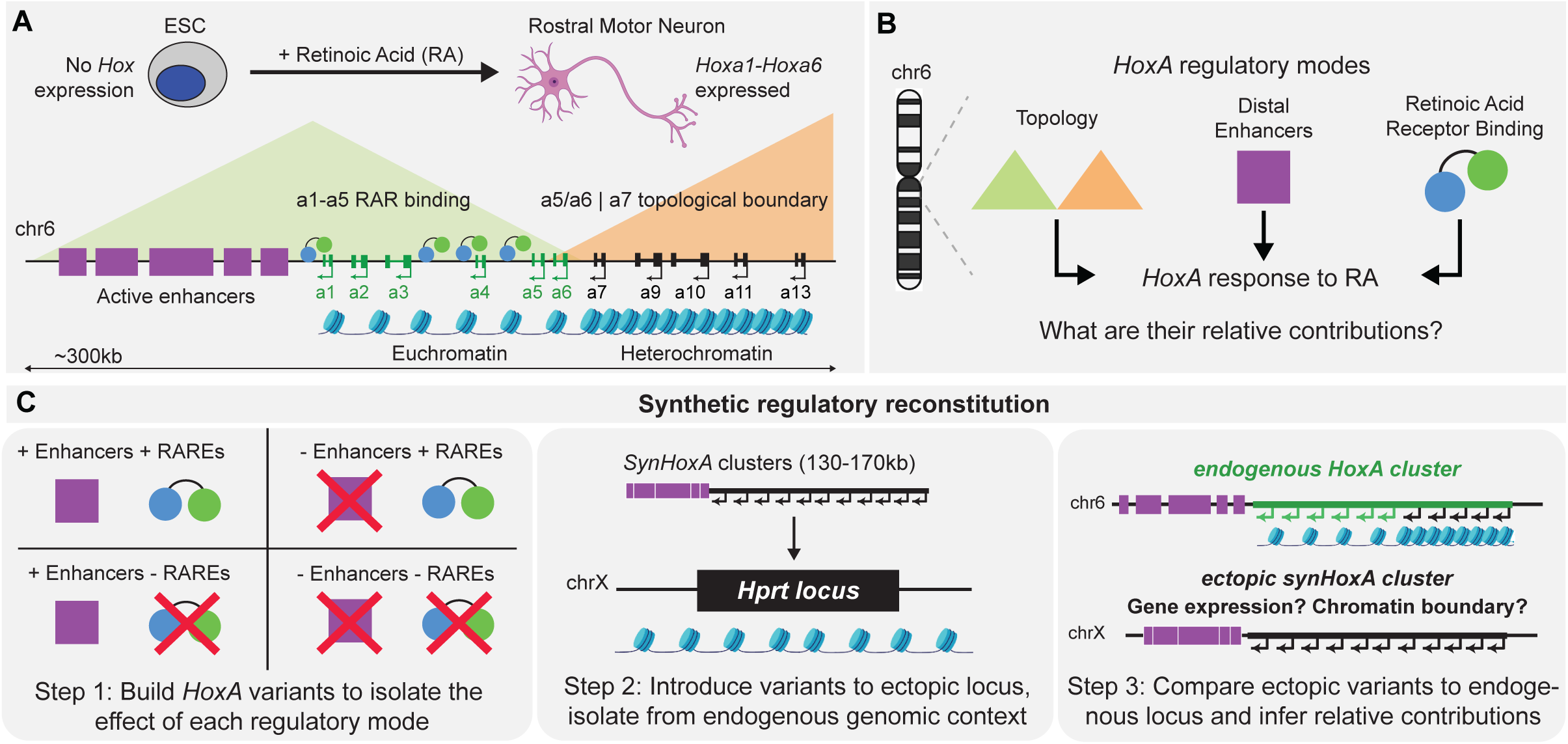
*HoxA* regulation relies on the integration of multiple regulatory modes. (A) Schematic of *HoxA* regulation in response to Retinoic Acid (RA). An *in vitro* mouse ES cell (ESC) – motor neuron differentiation model recapitulates *Hox* gene expression in response to RA. (B) Multiple regulatory modes, including enhancers, transcription factor binding and topology, are integrated to drive *HoxA* cluster response to RA. (C) Schematic of the synthetic regulatory reconstitution approach. Synthetic *HoxA* variants encoding various combinations of regulatory modes are built and integrated at an ectopic location in the genome. The response of these synthetic ectopic clusters to RA reveals their relative contribution to *HoxA* expression.

In undifferentiated cells, where no *Hox* genes are expressed, the entire *HoxA* cluster is targeted by the Polycomb Repressive Complex 2 (PRC2) (*18*) and is densely carpeted with the H3K27me3 histone mark, the PRC2 catalytic product. Patterning signals activate transcription through their downstream transcription factors and partition the *Hox* cluster into two chromatin domains (*19–21*). *Hox* clusters contain regulatory elements that bind these transcription factors (*22*). For example, the activated RA receptors (RAR) bind to RA responsive elements (RAREs) embedded within the *Hoxa1-Hoxa5* cluster domain (*19, 22, 23*). RA signaling leads to the separation of the cluster into two stable domains: active (H3K27ac marked), and inactive (H3K27me3 marked) chromatin, that contain transcribed and repressed genes, respectively (*19–21*) (Figure 1A).

The *Hox* cluster chromatin partition is mirrored by a topological reorganization in 3D, from a single globular domain into two domains harboring either transcribed or repressed genes (*20, 21, 24, 25*). In addition, the *HoxA* and *HoxD* clusters harbor topologically associated domain (TAD) boundaries (*12, 25, 26*) (Figure 1A). A strong topological boundary forms at CTCF binding sites within each of the clusters upon differentiation, separating the transcribed and repressed genes and promoting distal enhancer access to genes in the active domain (*14, 24, 25*). Deletion of CTCF motifs increases contacts between enhancers and repressed genes, and is associated with an expansion of the active chromatin domain, misexpression of *Hox* genes, and even homeotic transformations (*24, 25*).

In sum, distinct regulatory modes control *Hox* gene expression: local transcription factor binding, distal regulatory elements, and topological DNA organization. To date, it has been difficult to generate alleles that can separate the function of these modes. Therefore, a synergistic model describing their relative contribution and interactions has remained elusive. For example, is the endogenous genomic neighborhood and its organization required by CTCF and RAR to establish defined chromatin domains in response to RA? Are distal enhancers required for setting chromatin boundaries and the appropriate transcriptional output of single or multiple genes? Or do *Hox* clusters intrinsically contain the information required to activate the correct gene set in response to patterning signals (Figure 1B)?

Measuring the relative contributions of the different regulatory modes would require the generation of a set of ‘designer’ variant alleles over a sizeable genomic window. In addition, studying the direct effect of these genomic modifications on gene expression requires their isolation from confounding factors, such as the compensatory effect of other regulatory elements *in cis*. The ability to recapitulate endogenous regulation, including chromatin changes and transcriptional output, when placed at an ectopic genomic location is a very stringent test of the inherent regulatory potential of a DNA sequence. Therefore, attempting to reconstitute *HoxA* gene regulation at an ectopic locus would directly test the sufficiency of the transposed regulatory elements to drive various stages of *Hox* cluster regulation. Further, transposing variant *Hox* clusters containing subsets of regulatory elements enables the study of relative contributions of and interactions between the various regulatory modes.

Reconstituting fully editable *Hox* clusters has remained intractable primarily due to a lack of tools to precisely manipulate DNA at a scale that accurately models the complexity and size of native regulatory loci (>100kb) (*27*). While short reporter constructs enable the study of many variants over a small genomic window (<10kb), they suffer from a lack of controlled genomic context as they are largely randomly integrated or reside on episomal plasmids (*28*). Bacterial or Yeast Artificial Chromosomes (BACs/YACs) capture large genomic windows and have been shown to recapitulate endogenous gene expression after random transgenesis. However, they are not easy to manipulate, which makes it difficult to generate a large number of variants that can be tested *in vivo* (*29–31*). Further, methods for precise, single-copy integration of large DNA molecules in mammalian cells have been underutilized (*32, 33*). Recent advances in genome editing hold much promise for making precise modifications *in vivo*. However, it is still inefficient and time-consuming to make multiple defined edits in *cis*, phased on a single homolog (*34*). The focus has been on the impact of knockout/loss of function variants because gain of function and structural variants are less efficiently generated by CRISPR (*35*).

Recent advances in *de novo* DNA synthesis and assembly have enabled the construction of synthetic versions of the *Mycoplasma genitalium* (*36, 37*), *E. coli* (*38*), and yeast chromosomes (*39–45*). The final design of the synthetic yeast genome encodes a cluster of variants distributed on average every 400bp across 12 Mb of sequence, a feat that would be near impossible with top-down editing methods (*46*). The bottom-up assembly of loci enables the introduction of an arbitrary number of variants in *cis* that are independent of a natural template. Hence, variants such as complex structural rearrangements, multiplex editing, and insertion of novel DNA sequences across a large genomic window are tractable. We recently described a pipeline that harnesses the power of the endogenous homologous recombination machinery in yeast to *de novo* assemble ∼100kb regions of mammalian genomes and integrate them into a defined location in mouse embryonic stem cells (mESCs) (*47*).

Here we apply and extend this technology to the study of *Hox* cluster regulation. We *de novo* assembled variants of the rat *HoxA* cluster (ranging from 130 -170kb long) containing various combinations of the previously identified regulatory modes. We integrated them into a single-copy ectopic locus on the mouse X chromosome, thereby isolating them from the confounding effects of the native genomic neighborhood. We then asked whether variant ectopic clusters were sufficient to reconstitute the transcriptional and epigenetic *HoxA* cluster response to activating patterning signals (Figure 1C).

We found that a minimal cluster lacking enhancers and a cluster with all distal enhancer elements placed directly adjacent to *Hoxa1* drove the correct patterns of chromatin remodeling and transcription in response to the RA patterning signal, even in a foreign genomic neighborhood. Removal of the RAREs within the minimal *HoxA* cluster abolished all response to RA at the ectopic location. The introduction of enhancers to the *HoxA* cluster lacking RAREs was insufficient to fully rescue the loss of gene expression phenotype. This ‘synthetic regulatory reconstitution’ approach allows us to neatly separate the function of various regulatory modes and define the set of elements sufficient to drive various aspects of *Hox* gene regulation. We expect this will be a powerful approach to dissect the logic of transcriptional control in complex genomes.

## Results

We sought to understand the relative contributions of the various regulatory modes involved in the response of *HoxA* to patterning signals: local transcription factor binding sites, distal regulatory elements, and genomic neighborhood organization. The regulatory potential of a DNA sequence is best understood by testing its ability to drive gene expression predictably at an ectopic genomic location. Therefore, we tested the ability of *HoxA* cluster variants inserted at an ectopic genomic location to respond to patterning signals. We investigated the following unknowns: 1) role of genomic context in recruitment of repressing, activating, and boundary forming machinery during *HoxA* regulation; 2) distal enhancers’ quantitative contribution to *HoxA* regulation; and 3) role of intra-cluster regulatory elements.

To this end, we first relocated a synthetic cluster containing all elements previously described to be involved in endogenous *HoxA* regulation to an ectopic location on the X chromosome. To reveal the role of distal enhancers specifically, we synthesized a minimal construct lacking these sequences (Supplementary Figure 1A and Supplementary Figure 2A).

### Synthetic Hox strategy and construction

All constructs described here are derived from rat (*Rattus norvegicus*) *HoxA* cluster sequence, which shares ∼90% sequence similarity with the mouse sequence at *HoxA*. Although the sequence is highly conserved, polymorphisms enable distinction between ectopic and endogenous *HoxA* in sequencing-based analyses. Notably, this facilitated experiments in two distinct genetic backgrounds: 1) cells containing the endogenous *HoxA* cluster, which serves as an internal positive control for *Hox* regulation and for quantitatively comparing expression levels; and 2) cells lacking endogenous mouse *HoxA* to eliminate possible sequence mapping challenges.

We first constructed a 134kb wild-type rat minimal *HoxA* cluster (*SynHoxA*) by harnessing the homologous recombination machinery in yeast (*Saccharomyces cerevisiae*) (Figure 2 and Supplementary Figure 1). The minimal *SynHoxA* cluster contains all *HoxA* coding genes and encompasses the sequence corresponding to the repressed H3K27me3 contiguous domain in undifferentiated mESCs. We produced 28 ∼5kb PCR amplicons from BACs bearing the rat *HoxA* cluster, with ∼200 bp overlap between adjacent segments to enable homologous recombination. These amplicons were combined into ∼65kb rat *HoxA* half constructs upon co-transformation into yeast with appropriate linkers that direct assembly into an episomal yeast/*E. coli* shuttle vector. The half assemblies were recovered to *E. coli*, propagated, isolated, released from their BAC vectors and combined in yeast to produce the 134kb *SynHoxA* construct termed an “assemblon” (Supplementary Figure 1B). Edits to the assemblon can be made by switching the wild-type amplicons with synthetic DNA bearing the desired changes or by editing the assemblons directly using highly efficient, marker-free CRISPR Cas9-based engineering in yeast (*48*). This allows unfettered changes to pre-existing assemblons to be made rapidly (Figure 2).

**Figure 2:**
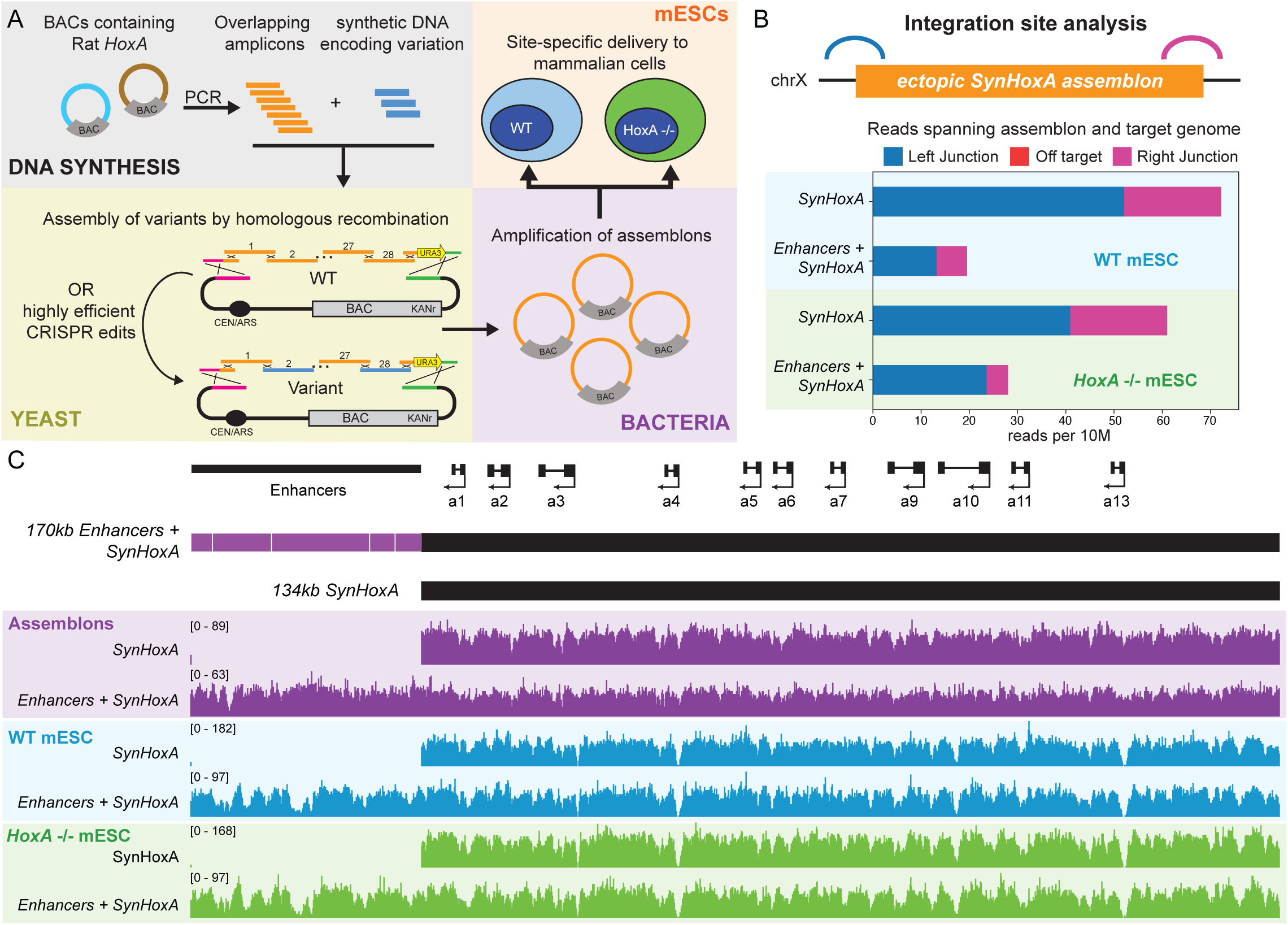
Build and delivery of *SynHoxA* constructs. (A) Schematic of the process to generate mESCs bearing ectopic synthetic *Hox* clusters via BACs containing Rat *HoxA*, overlapping PCR amplicons, homologous recombination-based assembly in yeast and amplification in bacteria. (B) Only the expected junctions spanning the synthetic assemblon and the host genome were observed in next generation sequencing data with no off-target integrations. (C) Schematic of the 134kb *SynHoxA* and 170kb *Enhancers+SynHoxA* assemblons, with the cluster in black and enhancers in purple along with the positions of protein coding genes. Sequencing data for assemblon DNA isolated from bacteria (purple) and from capture sequencing after integration in mESCs (blue and green). DNA sequencing data shown here are aligned to custom reference genomes (see Methods).

Several studies have previously identified regulatory elements that respond to RA (RARE elements) and/or Wnt (Ades for *Hox<u>A</u>* <u>d</u>evelopmental <u>e</u>arly <u>s</u>ide) in the gene-poor region distal to the endogenous *HoxA* cluster, between *Hoxa1* and *Skap2* (*14–16*). Deleting some of these regulatory regions reduces, but never eliminates, *HoxA* expression in response to RA or Wnt signaling (*14, 16*). To evaluate the importance of these enhancers, we generated a ‘compound enhancer’ that included all of these experimentally verified elements condensed together in ∼35 kb of DNA (Supplementary Figure 2). We fused this compound enhancer directly upstream of the 134kb core assemblon, resulting in a 170kb (*Enhancers+SynHoxA*) construct.

We performed DNA sequencing at each step of the pipeline (source BACs, half assemblons, and full assemblons) to confirm identity of the assemblon (Figure 2C and Supplementary Figure 3). Sequencing revealed that there were no gross abnormalities in the constructs. However, we find that there is a mutation frequency of about 1 single nucleotide polymorphism per 6kb on average arising from errors in PCR (Supplementary Figure 3). We verified that none of these mutations were likely to impact the characterization of these clusters in our differentiation system described below. The variants lie in intergenic space and are largely associated with relatively low conservation, except for a single exonic variant in *Hoxa7*, which is not expressed in mESCs or in response to RA. (Supplementary Table 1)

We used Inducible Cassette Exchange (ICE) for single-copy, site-specific delivery of the assemblons to the ectopic *Hprt* locus on the X chromosome of mESCs (*49*) (Supplementary Figure 5). *Hprt* is a housekeeping gene long used as a safe harbor site for genetic engineering studies (*50*). Although *Hprt* is in the middle of a TAD (26), recent work has shown remarkably little regulatory activity in the regions surrounding *Hprt* (*51*). For these reasons, we chose it as an appropriate neutral site to attempt regulatory reconstitution of the *HoxA* locus. All constructs described here were delivered to both *HoxA*^+/+^ and *HoxA*^-/-^ mESCs using ICE (Supplementary Figure 4). With ICE, on-target integration at *Hprt* was verified by phenotypic activation of conditional marker cassettes that depend on precise recombination events (G418^R^ and GFP^+^). Resistant mESC clones were validated by PCR with *SynHoxA* specific primers spanning the entire construct as well as novel junctions formed with the genome following integration (Supplementary Figure 5 E, F).

We used capture sequencing to validate that *SynHoxA* mESC clones contained the entire assemblon at *Hprt*. Nick-translation of our BACs yielded biotinylated probes specific to the synthetic ectopic locus, that were used to enrich for DNA from the assemblon, facilitating high-coverage sequencing at low cost (*52*). In all cases, we eliminated clones that contained deletions or rearrangements of *SynHoxA* sequence. (Figure 2C). mESC clones that passed the sequencing pipeline were further validated for precise on-target single-copy integration and absence of any off-target integration by analyzing reads spanning junctions between the synthetic construct and mouse genome (*52*). Importantly, we only detected reads spanning the expected junctions at the landing pad and no off-target integrations (Figure 2B). Therefore, we have established a powerful genome engineering technology at a scale that enables the study of transcriptional regulation at complex genomic loci, such as *Hox* clusters.

### Induction of SynHox expression during motor neuron differentiation

To investigate the response of the ectopic *SynHoxA* clusters to patterning signals, we chose a widely-used mESC differentiation system that recapitulates key aspects of ventral spinal cord development (*53–55*). After treatment with the ventralizing Hedgehog (Hh) and rostro-caudal RA patterning signals, mESCs transition through relevant progenitor states and differentiate into motor neurons (MNs) and interneurons (Figure 3A) (*53*). In this protocol, 90% of cells express *Hoxa5,* and have therefore collectively acquired an anterior or rostral identity (*56, 57*). Importantly, this differentiation strategy has provided critical insights into *Hox* regulation and insulating properties of CTCF that were ultimately confirmed *in vivo* (*19, 24, 25, 58*).

**Figure 3:**
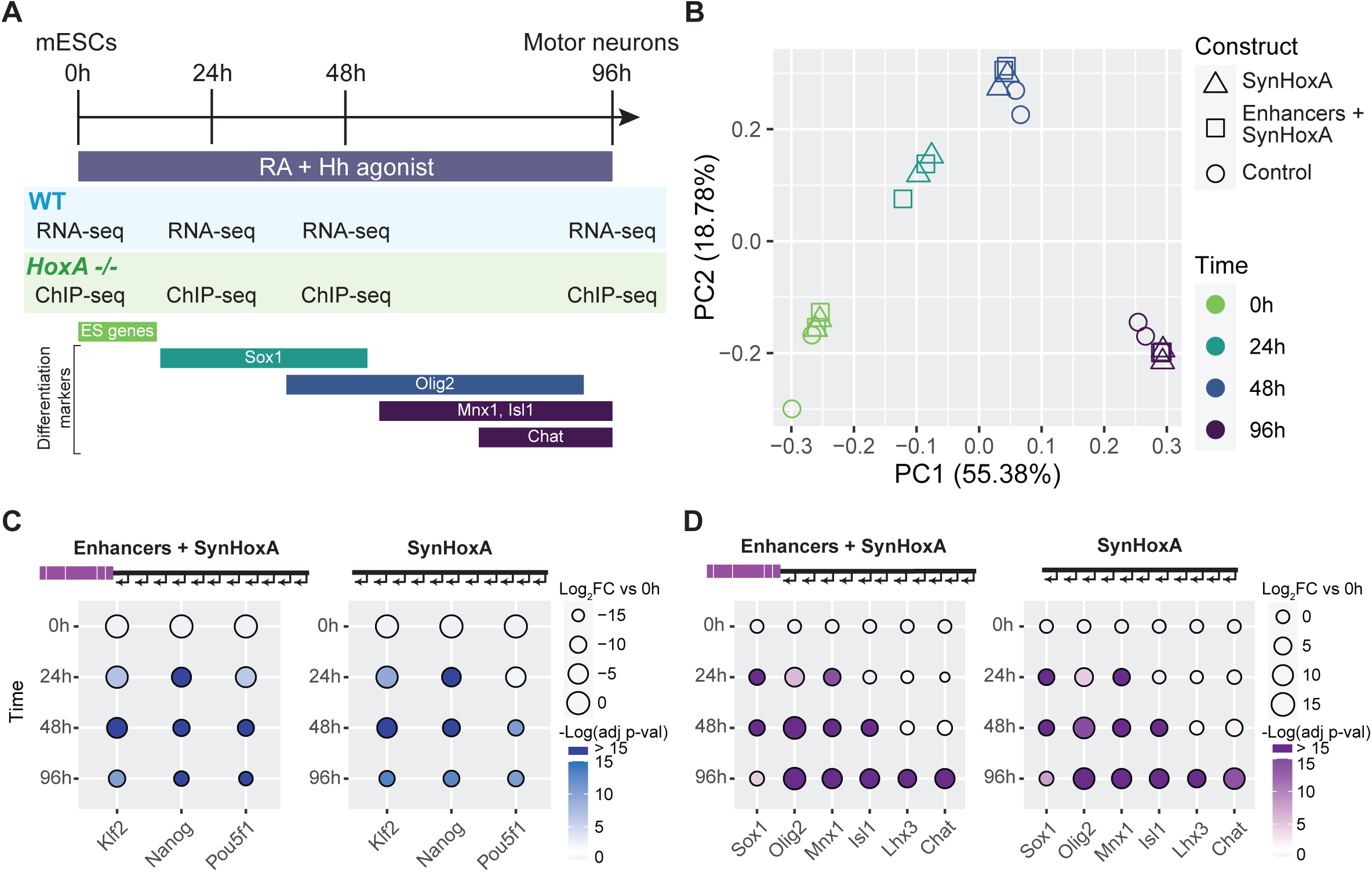
*SynHoxA* variants respond to patterning signals appropriately during *in vitro* spinal cord differentiation. (A) Overview of *in vitro* differentiation protocol. In response to RA and Hedgehog patterning signals, ESCs differentiate into MNs by transitioning through key progenitor states. WT and *HoxA* -/- mESCs harboring *SynHoxA* assemblons were analyzed by RNA-seq and ChIP-seq at indicated time points. Principal Component Analysis (PCA) of *SynHoxA* variant and control parental WT mESC lines RNA-seq datasets reveals clustering largely by time during the differentiation protocol (each data point represents independent differentiations). (C) Log2 fold change of pluripotency marker genes from RNA-seq data (n=2). *SynHoxA* variants downregulated pluripotency markers during differentiation as expected (n=2). (D) Log2 fold change of MN differentiation markers from RNA-seq data (n=2). *SynHoxA* variants upregulated differentiation markers as expected.

We investigated whether cells containing both endogenous *HoxA* and the *SynHoxA* clusters differentiate appropriately by performing RNA-seq over the course of the differentiation protocol, and by comparing to previously published control datasets (*59, 60*). Principal component analysis (PCA) revealed that the first two principal components explained 74% of the variance between datasets and that the samples grouped largely by time during the differentiation protocol (Figure 3B). Cells containing *SynHoxA* variant clusters downregulated pluripotency markers (Figure 3C), and upregulated markers of MN differentiation (Figure 3A, D). Thus, the integration of synthetic *Hox* clusters did not affect the ability of cells to differentiate appropriately after treatment with patterning signals.

### Distal enhancers are not required to specify active genes but boost early transcriptional output

*Hox* genes are repressed in their endogenous genomic context before exposure to patterning signals. In response to the ‘anterior’ signal RA, *Hoxa1-5* are induced at high levels, while the posterior *Hoxa7-13* remain repressed (*19*). We first asked whether the native configuration of distal enhancers and 3D organization are required for *Hox* clusters to appropriately activate genes in response to RA.

To that end, we differentiated cells with an intact endogenous *HoxA* cluster, additionally bearing the ectopic *Enhancers+SynHoxA* construct into MNs. As expected, the endogenous *HoxA* cluster induced the expected set of *Hoxa1-5* genes and had some weak *Hoxa6* expression at the latest time point as previously described (*19, 24*) (Figure 4A). Thus, the extra *SynHoxA* sequence did not affect endogenous *Hox* gene activation. The ectopic *Enhancers+SynHoxA* cluster induced *SynHoxa1*-*5* starting at 24h post RA (Figure 4B and Supplementary Figure 6). *SynHoxa4* and *SynHoxa5* mRNA levels increased as differentiation proceeded, and *SynHoxa6* to *SynHoxa13* remained repressed throughout. Therefore, the *Enhancers+SynHoxA* construct induced the correct subset of genes without misexpression of posterior genes during MN differentiation (Figure 4A, B and Supplementary Figure 6), showing that neither endogenous genomic context nor the wide spacing of enhancer elements is strictly required for a *Hox* cluster to induce the appropriate genes in response to a patterning signal.

**Figure 4:**
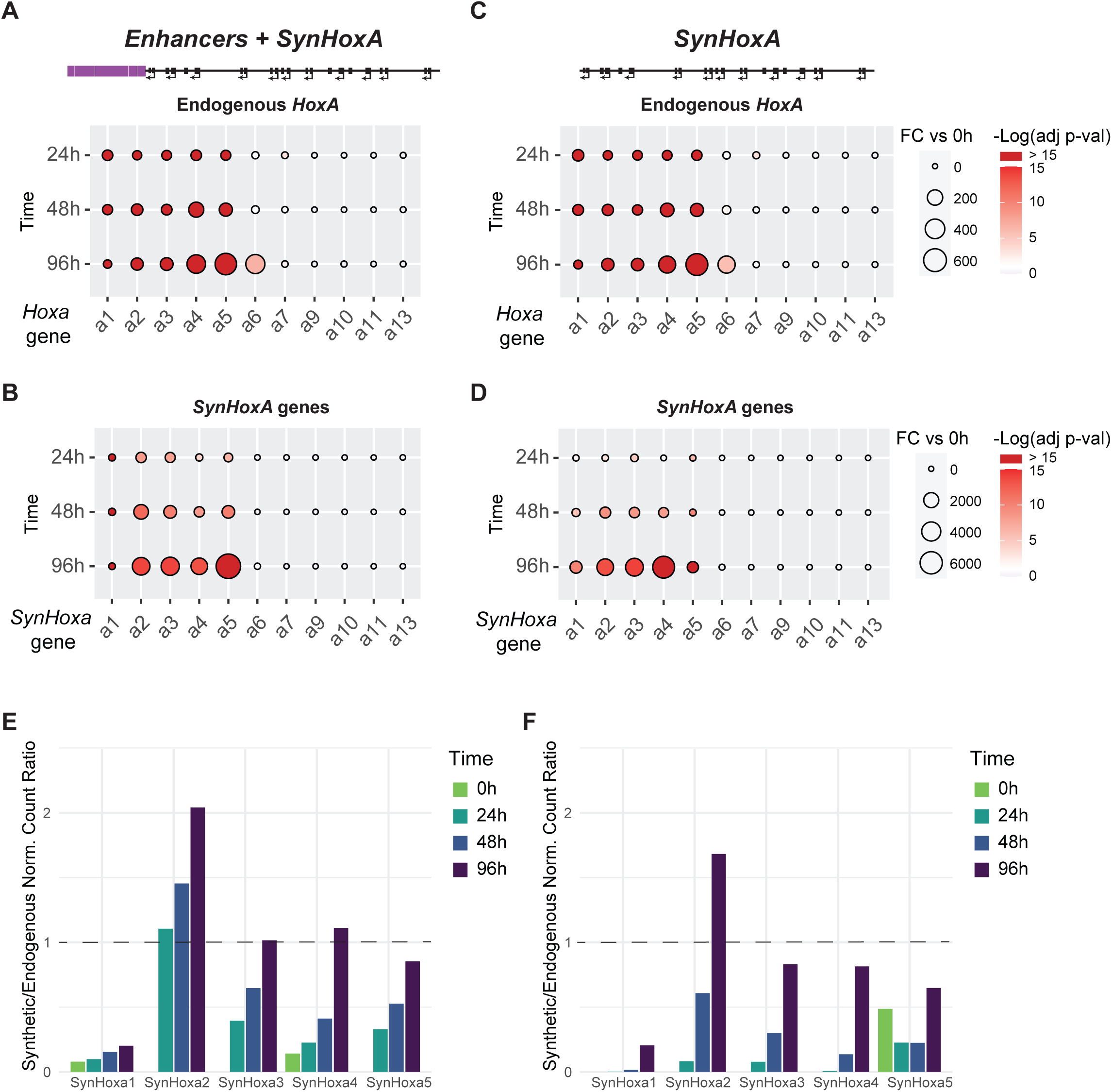
*SynHoxA* variants upregulate correct subset of *SynHoxA* genes in response to RA patterning signal. (A-D) Representation of RNA-seq data for endogenous mouse *HoxA* (A and C) and *SynHoxA* (B and D) genes during RA differentiation. Data are normalized to expression before RA treatment (0h) (n=2). *SynHoxA* variants upregulate the expected subset of genes (*SynHoxa1-5*) in response to the RA signal. (E-F) Ratios of gene expression for *SynHoxA* genes to endogenous mouse *HoxA* gene counterparts. (n=2). Counts for the endogenous *HoxA* genes were halved to normalize for two endogenous *HoxA* copies vs. one ectopic *SynHoxA* copy. RNA-seq data were aligned to a modified mm10 mouse genome containing the *SynHoxA* sequence inserted at the *Hprt* locus. Only uniquely-mapping reads were used in downstream analysis.

We sought to quantify the contribution of distal enhancers to *HoxA* gene activation. We repeated the differentiation experiment in the cell line containing the minimal *SynHoxA* and an intact endogenous *HoxA* cluster. Similar to the *Enhancers+SynHoxA* cluster, *SynHoxA* also specifically induced *SynHoxa1-5* (Figure 4C, D and Supplementary Figure 6). Therefore, the *HoxA* cluster itself contains all information that is required to decode the RA patterning signal into an anterior MN positional identity, independent of distal enhancers and the complex native genomic architecture.

We quantified *SynHoxA* gene induction by comparing each *SynHoxA* gene to its endogenous mouse *HoxA* counterpart (Figure 4E, F). There were some differences in the fine details of transcription from the *SynHox* clusters. Both *Enhancers+SynHoxA* and the minimal *SynHoxA* construct induced a lower amplitude of *SynHoxa1* transcription than the endogenous cluster. On the other hand, *SynHoxa2* induction surpassed the endogenous gene at 96h in both constructs. The induction kinetics of *SynHoxa3-5* were slower than *Hoxa3-5*, but the mRNA levels became comparable after 96h, particularly in the *Enhancers+SynHoxA* construct. Therefore, whereas a minimal *HoxA* cluster correctly responds to patterning signals, additional elements are required to fine-tune expression amplitude and timing.

The transcriptional response of the ectopic *HoxA* clusters can be summarized as follows. First, both constructs correctly upregulate *SynHoxa1-5* in response to anterior patterning signal RA during MN differentiation. Second, the addition of distal enhancers in *Enhancers+SynHoxA* leads to higher transcription levels, especially evident at earlier time points. Third, while the minimal *SynHoxA* is less effective at establishing high transcription levels early, the differences in mRNA levels lessen with differentiation time. Fourth, a direct comparison with endogenous genes revealed that while the most anterior *SynHoxa1-2* do not fully phenocopy endogenous expression levels, *SynHoxa3-5* highly resemble endogenous genes at the latest time point.

These data support a model in which the compact *Hox* clusters contain all necessary information to respond to extracellular signals by inducing the appropriate gene set. Distal enhancers are not required for specification of active genes but boost transcription and accelerate temporal dynamics. The endogenous genomic context and enhancer spacing may be required for further fine-tuning of early expression levels.

### Genomic context and distal enhancers are not required for PRC2 or CTCF recruitment

In ESCs, *Hox* clusters are carpeted with the repressive H3K27me3 histone modification (the PRC2 catalytic product) and recruit CTCF at potential boundary positions established in response to extracellular signals (*19, 24*). Active and repressive chromatin domains are established rapidly upon RA treatment (*19*). Therefore, we investigated whether the chromatin dynamics and CTCF recruitment of the relocated *HoxA* clusters mirrors their transcriptional output by performing ChIP-seq.

We used cells lacking the endogenous *HoxA* cluster to more reliably map reads to the *SynHoxA* clusters, thereby enhancing resolution. *SynHoxA* genes were expressed with similar dynamics in the presence or absence of the endogenous *HoxA* cluster (Supplementary Figure 7), indicating that *SynHoxA* regulation is primarily mediated in *cis*. Furthermore, the cells differentiated appropriately and acquire the expected MN fate (Supplementary Figure 11 A, C).

CTCF was recruited to similar sites within the ectopic *Enhancers+SynHoxA* and *SynHoxA* clusters as in the endogenous *HoxA* cluster (Figure 5A, B). Prior to exposure to patterning signals, the *Enhancers+SynHoxA* and *SynHoxA* clusters had high levels of H3K27me3 marks (Figure 5A, B). Thus, the *Hox* cluster, with or without distal enhancers, suffices to recruit the appropriate repressive chromatin marks and boundary elements even at an ectopic genomic location. The ability to recruit all components for correct patterning is intrinsic to the *HoxA* cluster sequence and is thus independent of endogenous genomic context or organization.

**Figure 5:**
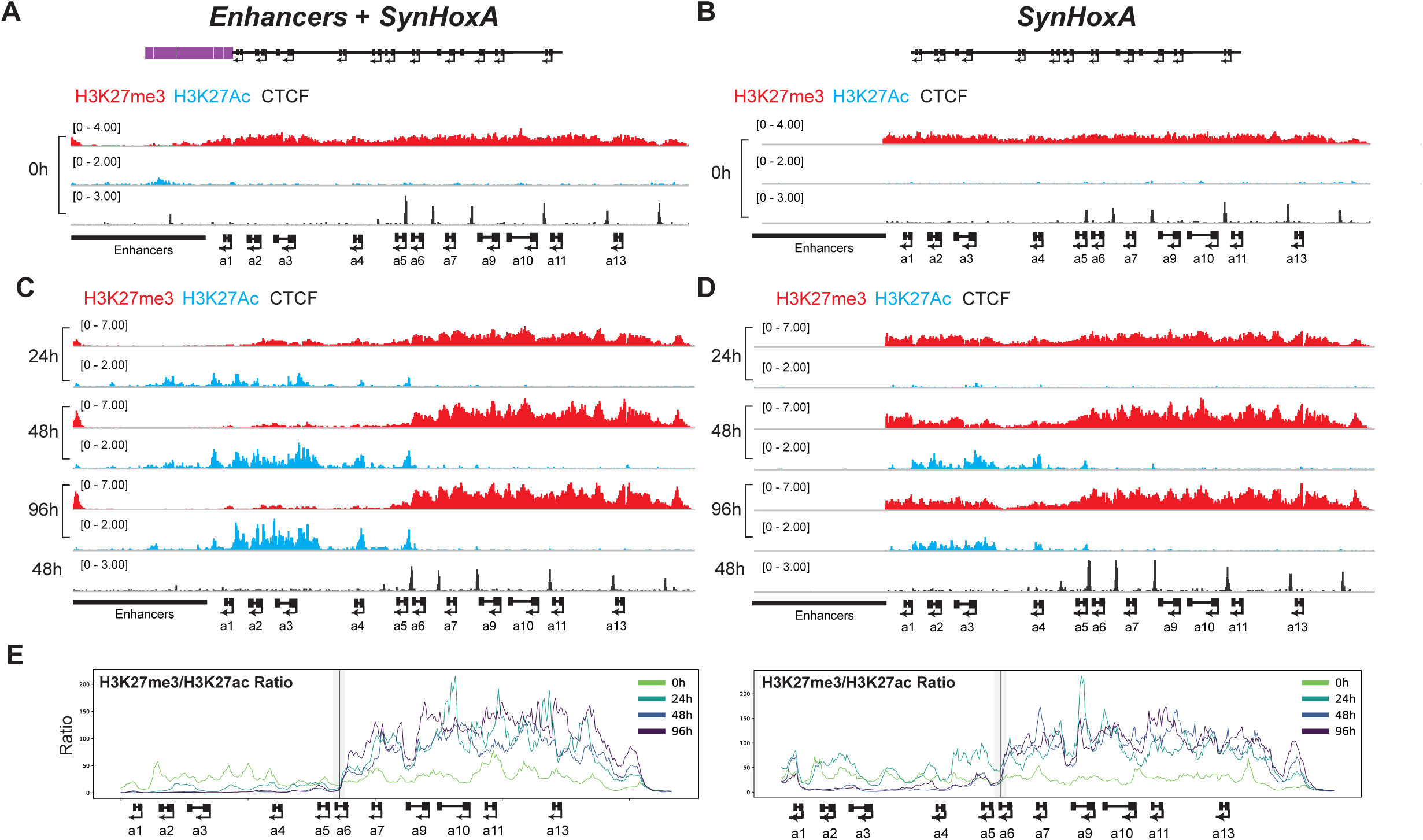
Distal enhancers are required for full clearance of repressive chromatin and formation of a sharp chromatin boundary. (A-B) Before exposure to patterning signals, *SynHoxA* clusters were decorated with repressive H3K27me3 chromatin marks (red), lacked activating chromatin H3K27Ac marks (blue), and recruited CTCF (black) to correct locations in ESCs (n=2). (C-D) In response to patterning signals, the *SynHoxa1-5* domain was marked by presence of activating H3K27Ac marks (blue) and clearance of repressive H3K27me3 marks (red). Presence of distal enhancers in the *Enhancers+SynHoxA* construct led to the formation of a sharper chromatin boundary. (E) Ratio of RPKM (Reads Per Kilobase per Million mapped reads) in 3kb windows sliding 300bp across *SynHoxA* of H3K27me3 and H3K27Ac ChIP-seq data from panels A-D reveals chromatin domain boundary between *Hoxa5* and *Hoxa6*. The black line marks the *Hoxa5* | *Hoxa6* CTCF site, with the extent of windows contributing to signal at the site shaded in gray. ChIP-seq data were aligned to a custom mm10 reference genome (see Methods).

### A minimal ectopic HoxA cluster establishes precise chromatin boundaries

The endogenous *HoxA* cluster responds to RA by forming a precise chromatin boundary between *Hoxa5* and *Hoxa6*, with some additional clearance of the repressive histone modification from *Hoxa6* at later time points (*19, 24*). The RA patterning signal leads to eviction of PRC2 from the *Hoxa1-5* domain with a concomitant H3K27ac increase, while maintaining PRC2 repression at *Hoxa7-13*, thus separating the *HoxA* cluster into two chromatin domains, active and inactive. We asked whether ectopic *Enhancers+SynHoxA* would partition into two chromatin domains during MN differentiation. Similar to the endogenous *HoxA* cluster, H3K27me3 at the *SynHoxa1-5* domain decreased while *SynHoxa6-13* gained H3K27me3 (Figure 5C). H3K27me3 removal from *SynHoxa1-5* was complemented by an increase in H3K27ac coverage, with no detectable H3K27ac deposited at *SynHoxa6-13*. H3K27me3 was entirely cleared by 48h, which is slightly slower than the endogenous locus, which clears by 24h (*19*). Therefore, the *Enhancers+SynHoxA* assemblon at an ectopic genomic location can respond to a patterning signal by forming two chromatin domains containing active and inactive *Hox* genes (Figure 5E) at exactly the correct positions. Thus, endogenous genomic context at *HoxA* is not required to translate an extracellular signal into an accurate epigenetic chromatin state.

We tested whether the minimal *SynHoxA* cluster would establish a chromatin boundary. Indeed, *SynHoxA* recruited H3K27ac at anterior *SynHoxA* genes upon RA activation (Figure 5D). Unlike the endogenous *HoxA* cluster and *Enhancers+SynHoxA,* H3K27me3 is not entirely removed from the *SynHoxa1-5* domain during differentiation (Figure 5D). Nonetheless, *SynHoxA* formed the appropriate, albeit weak, chromatin boundary at the *SynHoxa5-a6* CTCF binding site, evident from the H3K27me3 reduction at *SynHoxa1-5* (Figure 5E, Supplementary Figure 12).

These results suggest that the minimal *SynHoxA* cluster has the intrinsic ability to induce dynamic chromatin domains, independent of genomic context. This construct was unable to reduce H3K27me3 in the active domain to the same extent as the construct with enhancers. Either the boost in transcription provided by distal enhancers at early time points facilitates clearance of the repressive chromatin or the enhancers serve as platforms to recruit additional chromatin modifiers.

### Topological organization of ectopic SynHoxA clusters

The 3D structure of the *HoxA* cluster changes during MN differentiation, transitioning from a single association domain to two globular domains containing active or repressed chromatin (*24, 25*). Moreover, the active domain preferentially establishes associations with distal regulatory elements.

We investigated the topological organization of the *SynHoxA* clusters by performing Hi-C through differentiation (0 and 48h). Both ectopic *SynHoxA* clusters formed self-associating domains in undifferentiated cells without generating a *de novo* TAD boundary (Supplementary Figure 8). Similar to the endogenous cluster, *Enhancers+SynHoxA* broke into two domains during differentiation, with transcribed *SynHox* genes associating with enhancers in one domain and separating from inactive genes (Supplementary Figure 8A). The minimal *SynHoxA* similarly transitioned from a compact self-associated state in undifferentiated cells into two domains during differentiation (Supplementary Figure 8B). Taken together, the ectopic *HoxA* clusters have the intrinsic ability to self-organize in 3D, mirroring the expression and chromatin changes that occur upon differentiation at the endogenous cluster.

### The RARE sites within the HoxA cluster are required for the RA response

The minimal *SynHoxA,* containing all intra-cluster regulatory elements, transformed the RA signal into correct transcriptional and chromatin programs. This regulation could theoretically depend on either RAREs located within the *Hoxa1-5* domain, other sequences within the cluster, or it could rely on *de novo* long-distance regulatory interactions between the *SynHoxA* and surrounding regulatory elements. Therefore, we asked whether the RARE sites within *SynHoxA* are required for transcriptional activation and chromatin boundary formation. We built a third construct lacking all the RAREs (*RAREΔ SynHoxA*) and integrated it into WT and *HoxA*^-/-^ mESCs (Supplementary Figure 9 and Supplementary Figure 10). This assemblon had the following modifications from the minimal *SynHoxA*: 1) deletion of ∼3kb to remove the regulatory element located downstream of *SynHoxa1* and 2) precise mutations of the 3 other previously-described internal RARE sites (*22*).

Cells carrying *RAREΔ SynHoxA* differentiated appropriately, acquired the expected MN fate, and induced the appropriate endogenous *HoxA* genes (Figure 6A, Supplementary Figure 11B, D). However, unlike the *Enhancers+SynHoxA* and *SynHoxA* constructs, *RAREΔ SynHoxA* failed to upregulate *SynHoxa1-5* or form a chromatin boundary in response to RA signaling (Figure 6B-E, Supplementary Figure 12). *SynHoxa5* was the only *SynHoxA* gene with any signal at all, consisting of a few reads mapping to the assemblon. The explanation for this slight residual activity might lie in a nearby weakly conserved and poorly characterized RAR binding site located between *Hoxa5* and *Hoxa6* (*19, 22*).

**Figure 6:**
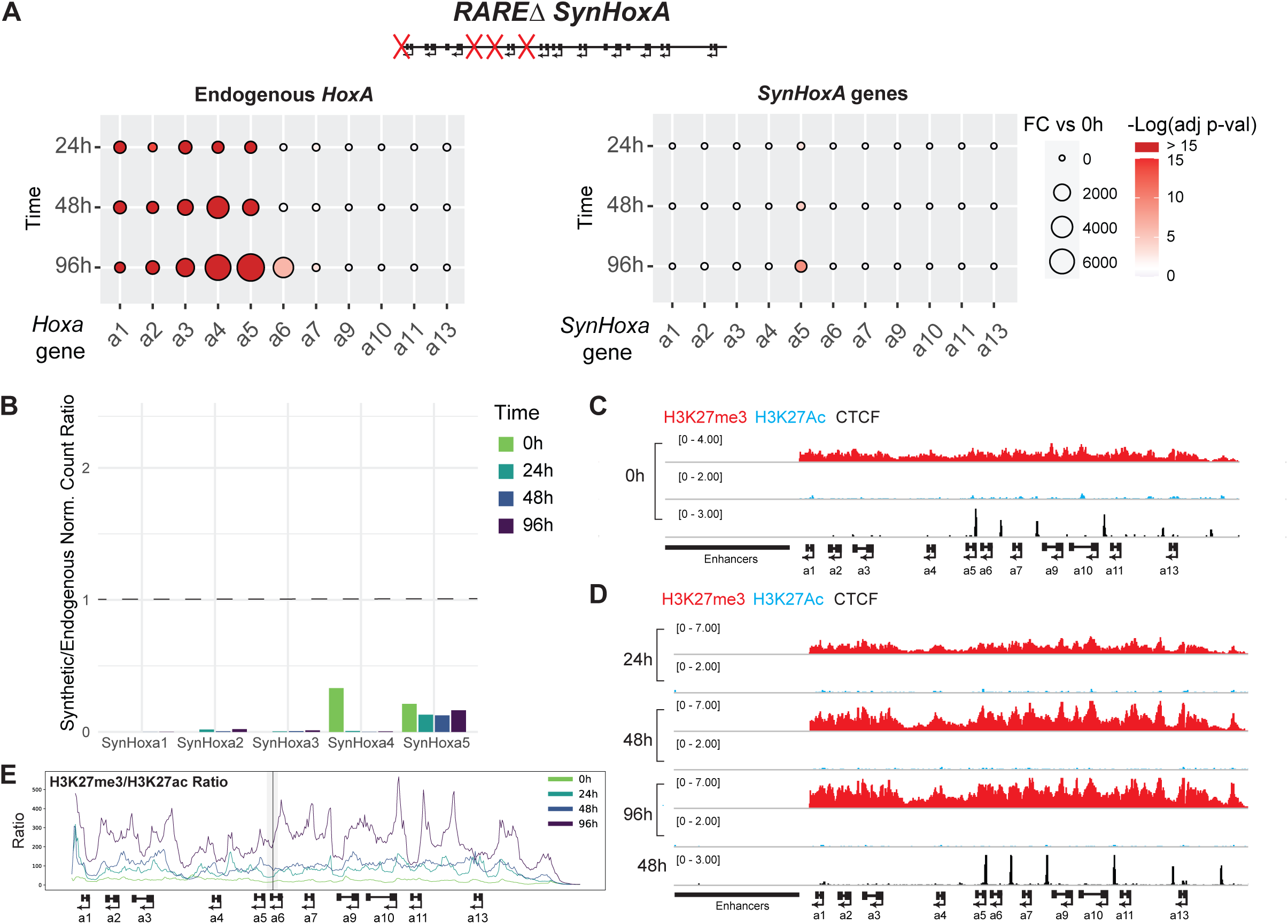
Retinoic acid receptor response element (RARE) sites are required for the RA response. (A) Expression fold-change of endogenous *HoxA* and *SynHoxA* genes during differentiation from RNA-seq data (n=2). *SynHoxA RAREΔ* did not upregulate *SynHoxa1-5* in response to RA signaling. Ratios of *SynHoxA* gene to endogenous *HoxA* gene expression from RNA-seq data (n=2), similar to Figure 4E,F. (C) Before exposure to patterning signals, *SynHoxA RAREΔ* was decorated with the repressive H3K27me3 chromatin mark (red), contained no evidence of the activating H3K27Ac chromatin mark (blue), and recruited CTCF (black) to the correct locations in ESCs (n=2). (D) There was no evidence of H3K27me3 (red) clearance and H3K27Ac (blue) recruitment at *SynHoxA RAREΔ*. CTCF (black) recruitment was unchanged. (E) No chromatin domain boundary was formed in response to RA at *SynHoxA RAREΔ.* Ratios of RPKM normalized repressive H3K27me3 to active H3K27Ac marks across *SynHoxA* of ChIP-seq data, similar to Figure 5E. The black line marks the *Hoxa5* | *Hoxa6* CTCF site, with the extent of windows contributing to signal at the site shaded in gray.

This lack of RA response provided an ideal background to measure the independent contribution of distal enhancers to *Hox* gene expression and chromatin state regulation. To that end, we built and integrated a fourth construct: *Enhancers + RAREΔ SynHoxA*. This assemblon contained all the enhancers inserted upstream of the *RAREΔ SynHoxA*. (Supplementary Figure 9 and Supplementary Figure 10). In *Enhancers + RAREΔ SynHoxA, SynHoxa2-5* were induced weakly in response to RA signaling, weaker than the minimal *SynHoxA* (compare Figure 7A-B with Figure 4). A weak chromatin boundary formed at the appropriate location between *SynHoxa5* and *SynHoxa6* (Figure 7C, D, Supplementary Figure 12). This suggests that distal enhancers have a weak ability to activate *HoxA* gene transcription independent of the internal RAREs’ driving force.

**Figure 7:**
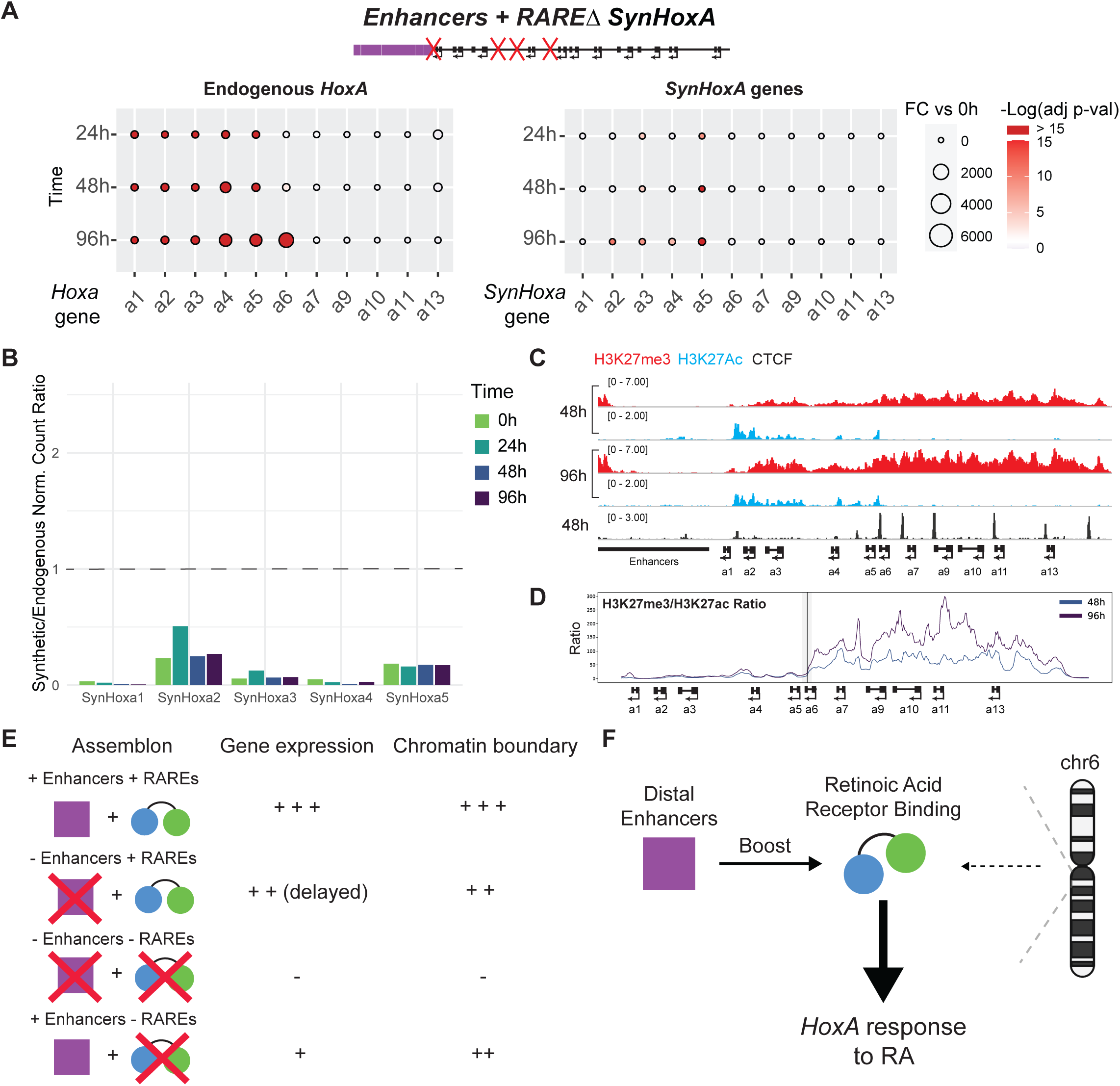
Addition of enhancers to *SynHoxA RAREΔ* does not rescue gene expression. (A) Fold change of *SynHoxA* or endogenous mouse *HoxA* genes during differentiation from RNA-seq data (n=2). Enhancers + *RAREΔ SynHoxA* weakly upregulated *SynHoxa1-5* in response to RA signaling. The endogenous *HoxA* cluster induced the correct set of genes. (B) Normalized ratios of gene expression for each *SynHoxA* gene to its endogenous mouse *HoxA* gene counterpart from the RNA-seq data (n=2). (C) Enhancers + *RAREΔ SynHoxA* formed a weak chromatin boundary, and recruited CTCF (black) to the correct locations in ESCs (n=2). (D) A chromatin domain boundary was formed in response to RA at the 5|6 CTCF site in Enhancers + *RAREΔ SynHoxA.* Ratios of RPKM normalized repressive H3K27me3 to active H3K27Ac chromatin marks from ChIP-seq data. Black line marks the *Hoxa5* | *Hoxa6* CTCF site, with the extent of windows contributing to signal at the site shaded in gray. (E) Summary of gene expression and chromatin boundary phenotypes across all *SynHoxA* clusters. (F) Model describing relative contributions of distal enhancers, intra-Hox RAR binding and genomic context to *HoxA* response to RA. RA signal is decoded by the RAR binding within the *HoxA* cluster. Distal enhancers increase the efficiency of chromatin remodeling and transcription. Endogenous genomic context is dispensable for *HoxA* cluster to respond to the RA signal.

## Discussion

Landmark discoveries on distal enhancer contribution to *Hox* gene expression relied primarily on deleting regulatory elements (*11, 14, 16*). Here we develop a ‘synthetic regulatory reconstitution’ approach and apply it to investigate the relative contributions of the regulatory modes (distal enhancers and intra-cluster transcription factor binding) that control *HoxA* gene expression. We systematically assembled and precisely integrated large DNA constructs (130-170kb) encoding *HoxA* variants into mESCs. We then differentiated these genetically modified pluripotent stem cells into motor neurons to study *Hox* gene expression and chromatin dynamics.

The *SynHox* cluster with enhancers (*Enhancers+SynHoxA*) induced the correct *Hox* gene set and formed a strong chromatin boundary at the correct location between *SynHoxa5* and *SynHoxa6.* The *SynHoxA* genes were expressed at levels comparable to the endogenous cluster at the latest time point. Some differences in gene expression dynamics were observed, suggesting that either specific regulatory elements or the endogenous genomic architecture and enhancer spacing are required for fine-tuning gene expression (*61*). Future work with larger transposed constructs that include the intervening sequences will help discriminate between these possibilities.

The minimal ectopic *SynHoxA* lacking distal enhancers recruited PRC2 and CTCF in embryonic stem cells and induced the correct subset of genes in response to RA. However, the dynamics of the response were delayed and reduced compared to the endogenous cluster and *Enhancers+SynHoxA*. This suggests that the distal enhancers’ primary effect is not to specify the set of genes that is active, but rather to boost transcription levels and chromatin turnover rate. These results are consistent with previous studies in which enhancers were deleted from the endogenous locus, leading to lower *Hox* gene transcription in response to RA or Wnt (*14, 16, 62*) and during development (*61, 63–65*).

We would like to highlight two relevant issues for PRC2 and CTCF recruitment. First, although canonical Polycomb Response Elements (PREs) for PRC recruitment have not been described in mammals, a small number of genomic locations recruit PRC2 that spreads to the rest of the PRC2-repressed genome based on linear or 3D proximity (*66*). These results suggest the *SynHoxA* clusters either have the intrinsic ability to recruit PRC2 or that they associate with distal recruitment sites, like the endogenous *Hox* clusters (*66*). Second, *HoxA* and *HoxD* clusters are at TAD boundaries (*12, 25, 26*). Therefore, CTCFs within the cluster engage in long-distance interactions with specific CTCF proteins bound at the other end of each TAD (*25*). Our results thus imply that precise CTCF positioning within the cluster does not depend on particular CTCF-CTCF interactions in the native locus or endogenous genomic organization.

Finally, we were able to study the relative contribution of distal enhancers and the regulatory elements that have been described within the *HoxA* cluster itself (*19, 22, 23*). This cluster lacking internal RAREs failed to respond at both the chromatin and transcriptional levels, supporting a model where *HoxA* internal RAREs are crucial for the RA response. Importantly, a cluster that contained distal enhancers but lacked RAREs did not fully rescue this phenotype, suggesting that internal RAR binding is the primary mode by which the RA signal is translated into an appropriate *HoxA* response. Distal enhancers serve to boost the dynamics of the transcriptional and chromatin response mediated by RAR binding upon receiving the RA differentiation signal. Altogether, we conclude that *Hox* clusters are regulatory units, with an intrinsic ability to respond to patterning signals, explaining in part the compact structure of vertebrate *Hox* clusters (Figure 7E, F).

This study is also the first proof-of-principle for a ‘synthetic regulatory reconstitution’ approach to understanding gene expression control. *In vitro* reconstitution has been a powerful framework to dissect complex biochemical processes because it allows for exquisite control over components of the system under study (*67, 68*). With this ability to generate locus-scale variant constructs that would have previously been intractable or have required numerous rounds of arduous gene editing, we expect synthetic regulatory reconstitution to revolutionize the study of transcriptional regulation.

## Acknowledgments

We want to thank all Mazzoni and Boeke lab members as well as the Institute for Systems Genetics community for their insightful comments and support. We thank Mohammed Khalfan, from the Genomics Core Facility at NYU, for making the reform tool for editing reference genomes publicly available; Bhavana Ragipani for preliminary observations and analysis on motor neuron differentiation markers; Natalie Zesati and Sonny Arora for help with preliminary visualization of the Hi-C data; Jane Skok and Danny Reinberg for insight and feedback. This work was supported in part by NHGRI RM1HG009491 to JB, MTM and EOM, NINDS R01NS100897 to EOM, and New York State Stem Cell Science pre-doctoral training grant C322560GG to MB.

## Author Contributions

SP, MB, EOM and JDB conceived of the project. SP, MB, DR, EH, RB and NE performed experiments. SP, MB, DR, LR, SM and MTM analyzed data. SG and JAC provided computational infrastructure support. MTM, LJH, EOM and JDB supervised research. SP, MB, EOM and JDB wrote the manuscript with input from all authors.

## Competing interests

JDB is a Founder and Director of CDI Labs, Inc., a Founder of and consultant to Neochromosome, Inc, a Founder, SAB member of and consultant to ReOpen Diagnostics, LLC and serves or served on the Scientific Advisory Board of the following: Sangamo, Inc., Modern Meadow, Inc., Sample6, Inc., Tessera Therapeutics, Inc. and the Wyss Institute. The other authors declare no competing interests.

## Data and materials availability

All sequencing data generated for this study will be deposited in NCBI GEO (pending).

## Materials and Methods

### Yeast and *E. coli* strains and media

All yeast work was performed in strain BY4741 using standard media and growth conditions. Yeast transformations were performed using the Lithium acetate method as previously described (*41, 69*). Plasmids were purified from bacterial cultures in Luria Broth (LB) supplemented with the appropriate antibiotic (12.5 µg/mL Chloramphenicol for source BACs, 25 µg/mL Kanamycin for all assemblons and 50 µg/mL Carbenicillin for all other plasmids). Plasmids purified from small scale bacterial cultures (5-10 mL) were used for all steps except for delivery to mESCs in which case DNA was purified from a large scale (500 mL) culture. Both bacteria and yeast were grown at 30°C.

A full list of yeast strains from this study is in Supplementary Table 2. A full list of plasmids from this study is in Supplementary Table 3.

### Yeast colony PCR

With the exception of the 134kb *SynHoxA* half-assemblon build (see relevant section in methods), all yeast colonies were genotyped by hand. Briefly, a single yeast colony was resuspended in 10-40 µl of 20 mM NaOH and placed in thermocycler. Yeast colony suspensions were boiled for 95°C for min and then cooled to 4C for at least 5 min before proceeding to PCR. 1ul of yeast lysate was used as template in a 10 µl GoTaq Green reaction (Promega M7123) with 0.25 µM of primers. PCR program: 95°C - 5min; 30X (95°C – 20s, 55°C – 90s, 72°C - 1min); 72°C – 5min; 4°C – hold. PCRs were separated on a 1% agarose gel containing ethidium bromide. 1kb Plus DNA Ladder (New England Biolabs N0550S) was used as a molecular weight standard.

### BAC recovery from yeast to *E. coli*

Plasmids were isolated from 5-10 mL yeast cultures either by alkaline lysis (*39*) or using Zymo Yeast Miniprep Kit I (Zymo Research D2001). Plasmids were then recovered into DH10B ElectroMax *E. coli* by electroporation (Invitrogen 18290015) following the manufacturer’s instructions.

### Yeast CRISPR/Cas9 editing

Donors bearing the desired mutations were generated by overlap extension PCR using Q5 polymerase (New England Biolabs M0492S) Given the high efficiency of homologous recombination following the generation of a defined double strand break, markers are not required to be encoded in the donor. This allows for seamless, marker-free editing. All CRISPR modifications in yeast were made as previously described (*70, 71*). Target strains were first transformed with the Cas9 plasmid (pNA519 pRS413- TEF1p-Cas9) and subsequently transformed with gRNA plasmids and linear donor fragments generated by PCR. Correct clones were identified by colony PCR followed by sequencing. Yeast CRISPR guide sequences are listed in Supplementary Table 3. Donor sequences are listed in the supplementary tables associated with each assemblon.

### Field Inversion Gel Electrophoresis (FIGE)

*SynHox* assemblon BACs were verified by digesting a ∼250-500ng purified by alkaline lysis (*72*) from small scale (5-10 mL) saturated bacterial culture with PvuI-HF (New England Biolabs R3150S). Digestion reactions were carried out at 37°C for 3-24 hours. The entire reaction was separated using the CHEF-DR system (Biorad 1703670) on a 1% low melting temperature agarose gel (Lonza 50100) in 0.5X TBE. The system was programmed using the auto algorithm function for fragments between 2kb and 50kb. 500 ng of lambda monocut ladder (New England Biolabs N3019S) were used as a molecular weight standard. Gels were stained after separation in a 0.5 µg/mL solution of ethidium bromide in water for 20-30 min and destained in water for 20-30 min before imaging.

### Design and build of 134kb *SynHoxA*

The 134kb *SynHoxA* assemblon was designed to cover the H3K27me3 domain at *HoxA* in mESCs (mm10 chr6:52151343-52285368). Corresponding rat coordinates (rn6 chr4:82120263-82339548) were identified using the UCSC genome browser Convert tool. The rn6 genome contains an erroneous duplication at *HoxA* between gaps in the assembly. We deleted this duplicated sequence *in silico* to arrive at the final 134kb *SynHoxA* sequence. Primers spanning the entire locus were designed using a custom script based on the Primer3 algorithm (*73*), yielding a list of 29628 unique 18-24bp primers that contain each of the 4 bases at a of 15% and meet cutoffs for primer dimer and hairpin formation scores. This list was then manually pruned to 28 primer pairs that cover the entire sequence with an average amplicon length of 4.5kb (range: 3.2kb-5.4kb) and an average overlap with the adjacent amplicon of 207bp (range: 363bp – 102bp). See Supplementary Table 4 for coordinates of design.

BACs containing Rat *HoxA* (CH230-79B14 and CH230-454G2) were obtained from the BACPAC resources center. DNA was isolated by alkaline lysis from 5-10 mL of overnight culture in LB supplemented with 12.5 µg/mL chloramphenicol. PCR amplicons were generated using Kapa HotStart Readymix (Kapa Biosystems KK2602) supplemented with 1 M Betaine (Sigma B0300-5VL) using 10-25 ng of BAC DNA as template and 0.3 uM of primers in a 20 µl reaction. All amplicons were generated with an annealing temperature of 68°C except for #12 and #16, which were generated at 65°C. PCR program: 95°C - 5min; 30X (98°C – 20s, 65°C/68°C – 30s, 72°C - 3min); 72°C – 5min; 4°C – hold.

134kb *SynHoxA* was built by first constructing two half-assemblons (segments 1-14 and segments 15-28). Two µl of each PCR amplicon were used to generate a pool, which was then transformed into yeast strain BY4741 with appropriate linkers (generated by overlap extension PCR) to direct assembly into a linearized vector (100 ng of I-SceI-digested pLM453) as previously described. (*47*) Linkers contained unique restriction enzyme sequences (I-SceI and AsiSI) to facilitate isolation of the insert sequence from the vector. The right linker for each of half assemblon also encoded a selectable *URA3* gene. In each assembly step, yeast transformants were first screened for the correct phenotypes (e.g. Ura+). Subsequently, colonies were tested for the presence/absence of assembly junctions using spanning primers (sequences are listed in Supplementary Table 5) across the construct with the aid of a robotic workcell as previously described (*47*). Plasmids from correct yeast clones (ySP0084 and ySP0085) were recovered into *E. coli* by electroporation to generate larger quantities of DNA.

To build the full 134kb *SynHoxA* assemblon, half-assemblon BACs (pSP0180 and pSP0182) were purified by alkaline lysis. Inserts were released using AsiSI digest and ∼1 µg of each were transformed into BY4741 with appropriate linker fragments and linearized vector (I-SceI digested pLM453). The right linker for this step encoded a *LEU2* marker, which allowed for the simple exclusion of yeast colonies arising from the transformation of undigested half-assemblon BACs (marked with *URA3*). Leu+ yeast colonies were screened for the presence of a PCR amplicon spanning segments 14-15 junction and for primers spanning the entire construct. Plasmids from correct yeast clones (ySP0088/89) were recovered to *E. coli* (pSP0193/196) and purified by alkaline lysis for verification by FIGE and sequencing.

In order to functionalize the assemblon for delivery by ICE, the backbone was modified in yeast using CRISPR. Strain bearing the full 134kb assemblon was transformed with Cas9 expressing plasmid pNA519 to yield yeast strain ySP0093. ySP0093 was subsequently transformed with a gRNA encoded in pSP0197 and a donor amplified from pLM707 containing the PGK-ATG-loxM-loxP-GFP cassette. The insertion was verified by Sanger sequencing. Plasmid was recovered from this yeast strain (ySP0096) into bacteria (pSP0211) and used for delivery to mESCs.

Supplementary Table 5 lists all sequences used in the build of 134kb *SynHoxA,* including segment primers, junction primers, linkers, gRNAs and reagents used for amplification of the donor used to insert the ICE cassette.

### Design and build of 170kb Enhancers + *SynHoxA*

Mouse enhancer coordinates were derived from published reports of enhancer function (*14–16*) and were expanded by adding 1 kb on each side. Mouse coordinates were then mapped to the rat genome using the UCSC genome browser Convert tool. In order to be conservative, we defined the final rat enhancer coordinates to be the union of those derived from the original and extended mouse enhancer coordinates (See Supplementary Table 6 for details). All enhancer sequences were then appended to yield the compound enhancer sequence. Primers were designed to amplify the requisite sequences as well as to generate linkers between non-contiguous segments (Supplementary Table 7).

PCR amplicons corresponding to the enhancer segments were generated using Q5 polymerase (New England Biolabs M0492S), except for segment 2 which was generated with Kapa polymerase supplemented with 1 M Betaine. 10-25 ng of CH230- 79B14 BAC prepped by alkaline lysis was used as template with 0.25 µM primers in 20 µl reactions. All amplicons were generated at an extension temperature of 68°C except for segment 2, which was generated at 72°C. PCR program: 95°C - 5min; 30X (98°C – 20s, 65°C – 30s, 68/72°C - 3min); 72°C – 5min; 4°C – hold.

PCR amplicons corresponding to the linkers between non-contiguous enhancer segments were generated by overlap extension PCR. Initial amplicons were generated with Q5 polymerase supplemented with 1 M Betaine. These were gel purified using Zymoclean Gel DNA Recovery Kit (Zymo Research D4002) and the two fragments were mixed together at a 1:1 ratio and 1-10 ng of this mix was used as template with outer primers in 20 µl Q5 reactions to generate the final linkers.

10 µl of each segment’s PCR product and linker product were pooled separately and then isopropanol-precipitated to a final volume of 10 µl of TE.

Yeast strain bearing the 134kb *SynHoxA* assemblon with delivery cassette (ySP0096) was transformed with Cas9 plasmid pNA519 to yield ySP0108. ySP0108 was transformed with plasmid pSP0233, which expresses a gRNA that cuts upstream of the 134kb assemblon, and with 10 µl of segment PCR pool and 5 µl of linker PCR pool to repair the gap. Yeast colonies were screened manually with junction primers spanning the overlaps between segments and with primers that span the 134kb *SynHoxA* assemblon (see Supplementary Table 7 for enhancer junction primers and Supplementary Table 12 for primers that span the 134kb *SynHoxA* assemblon).

A verified yeast colony (ySP0109) was recovered to bacteria (pSP0242) for verification by FIGE and sequencing.

### Design and build of 130kb *RAREΔ SynHoxA* and 166kb *Enhancers* + *RAREΔ SynHoxA*

Rat RARE sequences were based on published reports of mouse RAREs that were converted to Rat coordinates using the UCSC genome browser Convert tool (*22*). The region containing 3’Hoxa1 RARE (rn6 chr4:82120263-82123922) was deleted. Other RARE mutations were designed such that only the direct repeat sequences were mutated to polyA. Mutagenesis was performed using CRISPR/Cas9. Donors were generated by overlap extension PCR (Supplementary Table 8).

Hoxa7 CDS mutation R131W was corrected in ySP0108 (yeast strain with 134kb *SynHoxA –* ICE delivery cassette + pNA519 Cas9 plasmid) by CRISPR/Cas9. ySP0108 was transformed with gRNA expression plasmid pSP0255 and a donor generated from a synthetic oligo obtained from IDT (Supplementary Table 8). Yeast colonies were screened by PCR followed by Sanger sequencing to yield wild type Hoxa7 strain ySP0126, which was transformed with Cas9 expression plasmid pNA519 to yield ySP0128, which is the parent to all subsequent assemblons.

3’Hoxa1 RARE was first deleted using gRNA expression plasmid pSP0277 to yield ySP0131. ySP0131 was transformed with pNA519 Cas9 expression plasmid to yield ySP0146. 5’Hoxa3 RARE and 5’Hoxa4 RAREs were modified in ySP0146 with gRNAs expressed from pSP0322 and requisite donors to yield ySP0147. Finally, 3’Hoxa4 RARE was mutated in ySP0147 using the gRNA expression plasmid pSP0315 to yield ySP0161. All mutations were verified in yeast colonies using genotyping primers specific to the mutation (Supplementary Table 8) followed by Sanger sequencing of the region,

To add enhancer sequences to the 130kb *RAREΔ SynHoxA* assemblon, enhancers were amplified from the 170kb *Enhancers + SynHoxA* BAC (see section on build of 170kb assemblon). A pool of enhancer amplicons was transformed into ySP0161 with gRNA expression plasmid pSP0233 and linkers to repair the cut upstream of the assemblon. Yeast colonies were screened for the presence of all enhancer junctions as well as sequences spanning the 134kb S*ynHoxA* assembly (see Supplementary Table 7 for enhancer junction primers and Supplementary Table 12 for primers that span the 134kb *SynHoxA* assemblon) The plasmids from ySP0161 and ySP0200 were recovered to bacteria and subject to verification by FIGE and sequencing to yield pSP0328 and pSP0394, respectively.

### mESC culture, media and differentiation

A17iCre mESCs (*49*) were cultured on plastic tissue culture plates coated with 0.1% gelatin (EMD Millipore ES-006-B) at 37°C and 5% CO2 in ‘80/20’ medium comprising 80% 2i medium and 20% mESC medium. mESCs were grown at 37°C and 8% CO2 before differentiation experiments.

2i medium was made from a 1:1 mix of Advanced DMEM/F12 (Gibco 12634010) and Neurobasal-A (Gibco 10888022), containing 1X N-2 supplement (Gibco 17502048), 1X B-27 supplement (Gibco 17504044), 2 mM Glutamax (Gibco 35050061), 0.1 mM Beta-Mercaptoethanol (Gibco 31350010), 10^3^ units/mL LIF (Millipore, ESG1107), 1 μM MEK1/2 inhibitor (Stemgent, PD0325901), and 3 μM GSK3 inhibitor (R&D Systems, CHIR99021). mESC medium was made from Knockout DMEM (Gibco 10829018), containing 15% Fetal Bovine Serum (Gemini), 0.1 mM Beta-Mercaptoethanol (Gibco 31350010), 1X MEM Non Essential Amino Acids (Gibco 11140050), Glutamax (Gibco 35050061), 1X Nucleosides (Millipore ES-008-D) and LIF (Millipore ESG1107).

The protocol for mESC *in vitro* differentiation to motor neurons has been described previously (*53*). Briefly, trypsinized (Gibco) mESCs were plated in AK medium (Advanced DMEM/F12:Neurobasal (1:1) medium (Gibco), 7% KnockOut SR (vol/vol) (Gibco), 2 mM L-glutamine, 0.1 mM ß-mercaptoethanol and penicillin–streptomycin (Gibco)) to induce formation of embryoid bodies (EBs), at 37°C, 5% CO2. 3.5×10^5^ cells were plated in 100 mm suspension plates (Corning) for RNA-seq experiments, while 3.5×10^6^ cells were plated in 245 mm x 245 mm square plates (Corning) for ChIP-seq experiments. After 2 days (0h timepoint), the EBs were split 1:2 and plated in fresh AK medium supplemented with 1 μM all-trans retinoic acid (RA) and 0.5 μM smoothened agonist (SAG) (Millipore 566660).

Supplementary Table 9 lists all cell lines used in this study.

### PCR genotyping of mESCs

mESC clones were initially screened by performing PCR on crude gDNA extracted from cells growing in a 96-well plate. Briefly, cells were washed once with PBS and frozen at -80°C for at least 30 min. The plate was thawed at room temperature and cells were resuspended in 40-50 µl of TE supplemented with 0.3 µg/ul Proteinase K (Thermo Scientific EO0492) and transferred to a 96-well PCR plate. Cell suspensions were then incubated in a thermocycler at 37°C for 1 hour and then 99°C for 10 min. 1 µl of this crude lysate was used for PCR in a 10 µl GoTaq Green reaction (Promega M7123) with 0.25 µM of primers. Candidate clones were expanded and re-genotyped using DNA extracted with the QIAamp DNA mini kit (Qiagen 51306) according to the manufacturer’s protocol. 25-50 ng of DNA was used as template per reaction. PCRs were separated on a 1% agarose gel containing ethidium bromide. 1kb Plus DNA Ladder (New England Biolabs N0550S) was used as a molecular weight standard.

### Generation of *HoxA* ^-/-^ cell line

Deletion of the endogenous *HoxA* alleles was designed to mimic the sequence of 134kb *SynHoxA* (mm10 chr6:52151380-52285416). gRNAs were designed using CRISPOR and were chosen to have high cutting efficiency and low off-target scores (*74*). gRNAs were cloned into pSP0172, a modified version of pX459 (Addgene #62988, a gift from Feng Zhang) that has the Puromycin resistance gene replaced by a Blasticidin resistance gene. A 200 bp single stranded oligo donor (ssODN, oSP379) was designed to bridge the deletion and ordered from Integrated DNA Technologies (IDT).

gRNA expression plasmids were purified using PureLink HiPure Plasmid Midiprep Kit (Invitrogen K210004) following manufacturer’s instructions. 12.5 µg of gRNA expression plasmids pSP0161 and pSP0164 were nucleofected into 1 million A17iCre mESCs along with 5 µl of 100 µM ssODN. Cells were transiently selected with 10µg/mL Blasticidin (Invivogen ant-bl-05b) for 3 days post nucleofection. Single clones were picked and genotyped using primers that span the deletion junctions and with primers internal to mouse *HoxA*. Sanger sequencing confirmed the precise deletion. Clones with the correct genotype were expanded and subject to WGS. Clones were also verified by metaphase spread karyotyping and quantitative real-time PCR (qRT-PCR) for ESC markers as previously described (*47*). A passing clone (1-A6) was used for all experiments.

Sequences of guides, donor and genotyping primers can be found in Supplementary Table 10. Sequences of qRT-PCR primers are in Supplementary Table 11.

### Delivery of assemblons to mESCs and verification

Bacterial strain carrying the desired assemblon was struck out and grown for 2 days at 30°C on LB-agar plates containing 50 µg/mL Kanamycin. Single colonies were picked into 5 mL of LB + 25 µg/mL Kanamycin and grown for approximately 8 hours. The starter culture was used to seed 500 mL cultures at a 1:1000 dilution. 500 mL cultures were shaken at 30°C for 18-24 hours. Cells were pelleted and left at -20°C for long term storage or directly used for isolating DNA. BAC DNA was isolated using Nucleobond XtraBAC kit (Takara 740436.25) according to manufacturer’s protocol. DNA pellets from 500 mL cultures were resuspended in 40 µl of TE.

DNA was delivered by nucleofection using the Nucleofector 2b system (Lonza VPH-1001). 1.5 million A17iCre mESCs were plated on gelatin-coated 10 cm dishes 2 days before delivery. Cre expression was induced with 1 µg/mL doxycycline 18-24 hours before delivery. On the day of delivery, cells were washed with PBS and harvested using Accutase (Biolegene 423201). 5-6 million cells were used per delivery. 10 µl purified assemblon DNA were used per delivery. Cells were resuspended in 100 µl nucleofection solution with purified DNA and transferred to a cuvette using wide bore tips. Cells were nucleofected using program A-023 and plated on gelatinized 10 cm dishes. Cells were selected with 400 µg/mL Geneticin (Life Technologies, 10131-027) 48 hours post nucleofection. Resistant clones were picked 10-14 days post nucleofection.

Crude gDNA was extracted from clones and PCR genotyping was performed with primers spanning genome junctions (tetO-GFP and PGK-Neo), primers specific to endogenous *HoxA* deletion and primers specific to the overwritten *Cre* in the landing pad. In addition, clones were screened with a subset of heterologous primers that were designed using Primer-Blast to be specific to *SynHoxA* sequences when compared to endogenous mouse *HoxA*. Correct clones were expanded, pure genomic DNA was extracted and clones were genotyped with the full complement of primers. Passing clones were then verified to contain the full assemblon by capture sequencing. Sequences of genotyping primers are in Supplementary Table 12.

To generate the flow cytometry plot in Supp. Fig 5, parental WT mESCs and capture-seq verified WT mESCs bearing the 134kb *SynHoxA* assemblon at *Hprt* were treated with 3 µg/mL doxycycline for 24 hours. Flow cytometry was performed on a BD Accuri C6 instrument and results were analyzed using FlowJo. Cells were gated on forward and side-scatter and histograms of GFP expression normalized to mode were plotted.

### Library preparation for next generation sequencing

A list of all sequencing libraries and information associated with them can be found in Supplementary Table 13.

#### BAC DNA sequencing

Illumina sequencing libraries were generated from 100-200 ng BAC DNA using NEBNext Ultra II FS DNA Library Prep Kit (New England Biolabs E7805S) according to the manufacturer’s protocol. Libraries were sequenced on an Illumina NextSeq 500 in paired end mode.

#### Sequencing mESCs (WGS and Capture-Seq)

mESC gDNA was purified from 1-5 million cells using the QIAamp DNA mini kit (Qiagen 51306) according to the manufacturer’s protocol. Illumina libraries were prepared as previously described (*52*). 1 µg of DNA was sheared to ∼500 to 900 bp in a 96-well microplate using the Covaris LE220 (450 W, 10% Duty Factor, 200 cycles per burst, and 90-s treatment time). Sheared DNA was purified using the DNA Clean and Concentrate-5 Kit (Zymo Research D4013), and the concentration was measured on a NanoDrop instrument (Invitrogen). DNA fragments were end-repaired with T4 DNA polymerase, Klenow DNA polymerase, and T4 polynucleotide kinase (New England Biolabs M0203S, M0210S and M0201S, respectively), and A-tailed using Klenow (3′-5′ exo-; New England Biolabs M0212L). Illumina-compatible adapters were subsequently ligated to DNA ends, and DNA libraries were amplified with KAPA 2X Hi-Fi Hotstart Readymix (Roche).

Libraries for whole genome sequencing (WGS) of parental mESC lines (WT and *HoxA*^-/-^ ) were sequenced on a Novaseq 6000 in paired end mode.

Targeted sequencing using in-solution hybridization capture (Capture-seq) was performed on *SynHoxA-*delivered mESCs as previously described (*52*). Biotinylated baits for capture sequencing were prepared from assemblon BACs using nick translation. 134kb *SynHoxA* mESCs were captured with bait made from 134kb *SynHoxA* BAC. All other mESCs were captured with bait made from 170kb *Enhancers* + *SynHoxA* BAC. In addition, the parental mESCs and RAREΔ *SynHoxA* mESCs were captured with a bait that included the ICE landing pad and flanking mouse genome sequence (pLM1103+ICEFlanking). See Supplementary Table 13 for details. All libraries from *SynHoxA* deliveries were sequenced on an Illumina NextSeq 500 in paired end mode.

#### ChIP-seq

Cells were collected at 0h or 24h, 48h, or 96h after RA/SAG treatment. ChIP-seq was performed as previously described (*75*).

Cells were crosslinked at room temperature in 1 mM DSG (ProteoChem) for 15 min, followed by the addition of 1% FA (vol/vol) for 15 min. The reaction was quenched with Glycine and cells were washed with 1 × PBS. Samples were divided into ∼25-30 million cell aliquots, pelleted by centrifugation at 275 g, and frozen at -80°C. Cell aliquots were thawed on ice and lysis was performed in 5 mL of 50 mM HEPES-KOH pH 7.5, 140 mM NaCl, 1 mM EDTA pH 8.0, 10% glycerol (vol/vol), 0.5% Igepal (vol/vol), 0.25% Triton X- 100 (vol/vol) with 1 × protease inhibitors (Roche, 11697498001) for 10 min at 4°C. Cells were pelleted by centrifugation for 5 min at 1200 g, resuspended in 5 mL of 10 mM Tris-HCl pH 8.0, 200 mM NaCl, 1 mM EDTA pH 8.0, 0.5 mM EGTA pH 8.0 with 1 × protease inhibitors, and incubated for 10 min at 4°C on a rotating platform. Cells were centrifuged for 5 min at 1200 g and resuspended in 2 mL of Sonication Buffer (50 mM Hepes pH 7.5, 140 mM NaCl, 1 mM EDTA pH 8.0, 1 mM EGTA pH 8.0, 1% Triton X-100 (vol/vol), 0.1% sodium deoxycholate (wt/vol), 0.1% SDS (vol/vol) with 1 × protease inhibitors). For sonication, each sample was split in two Bioruptor tubes with added sonication beads. Sonication was performed using the Bioruptor Pico (Diagenode) for 18 cycles of 30 sec on and 30 sec off to sheer DNA into an average size of approximately 200bp. Immunoprecipitation was performed for 16h at 4°C on a rotating platform by incubating with Dynabeads protein-G (Thermo Fisher Scientific) conjugated with antibodies. For histone modifications, half of each original cell aliquot was incubated with Dynabeads protein-G conjugated with 5 µg of rabbit polyclonal to Histone H3K27me3 (Active motif 39155) or rabbit polyclonal to Histone H3 (acetyl K27) (Abcam ab4729) antibodies. For CTCF, the entire cell aliquot was incubated with Dynabeads protein-G conjugated with 5 µl of rabbit polyclonal to CTCF (EMD 07-729) antibody.

After the immunoprecipitation, washes were performed with the following ice-cold buffers: sonication buffer, sonication buffer with 500 mM NaCl, LiCl wash buffer (20 mM Tris-HCl pH 8.0, 1 mM EDTA pH 8.0, 250 mM LiCl, 0.5% Igepal (vol/vol), 0.5% sodium deoxycholate (wt/vol)), and TE buffer (10 mM Tris-HCl pH 8.0, 1 mM EDTA pH 8). Elution was performed in elution buffer (50 mM Tris-HCl pH 8.0, 10 mM EDTA pH 8.0, 1% SDS (vol/vol)) by incubating for 45 min at 65°C with occasional flicking of the tube. Samples were incubating for 16h at 65°C to perform reversal of crosslinks. 200 μL of TE and RNase A (Sigma) at a final concentration of 0.2 mg/mL was added to digest RNA by and incubating for 2h at 37°C. Proteins were digested by adding Proteinase K (Invitrogen) at a final concentration of 0.2 mg/mL, supplemented with CaCl2, at 55°C for 30 min. DNA was purified with phenol:chloroform:isoamyl alcohol (25:24:1; vol/vol) (Invitrogen) followed by an ethanol precipitation. DNA pellets were resuspended in 70 μL of water. lllumina DNA sequencing libraries were prepared with approximately one third of the ChIP sample (24 μL) or a 1:100 dilution of the input sample in water. Library preparation was performed by end repair, A-tailing and ligating Illumina-compatible Bioo Scientific multiplexed adapters. Agencourt AmpureXP beads (Beckman Coulter) were used to remove unligated adapters. PCR amplification was performed with Phusion polymerase (New England Biolabs) and TruSeq primers (Sigma). Libraries were gel purified (Qiagen) between 250 and 550bp in size. Libraries were quantified before pooling using the KAPA library amplification kit on the Roche Lightcycler 480 or the Bio-Rad CFX96. The libraries were sequenced on Illumina NextSeq 500 using V2.5 chemistry (75 cycles, single-end 75bp) or on Illumina NovaSeq 6000 using the SP Reagent Kit (100 cycles, single-end 100bp) at the Genomics Core Facility at NYU.

#### RNA-seq

Cells were collected at 0h or 24h, 48h, or 96h after RA/SAG treatment. RNA-seq was performed as previously described(*75*). RNA was extracted with the TRIzol LS Reagent (Life Technologies) followed by purification with the RNAeasy mini kit (Qiagen). RNA integrity was checked with the Agilent High Sensitivity RNA Screentape (Agilent, 5067-5579). 500 ng of RNA was used to prepare libraries and spiked-in with ERCC Exfold Spike-in mixes (Thermo Fisher 4456739). The TruSeq Stranded mRNA Library Preparation kit (Illumina 20020594) was used to prepare RNA-seq libraries. Library size was checked on the High Sensitivity DNA ScreenTape (Agilent 5067-5584). The KAPA library amplification kit was used to quantify libraries on the Bio-Rad CFX96 or the Roche Lightcycler 480 before pooling libraries. Libraries were sequenced on the Illumina NextSeq 500 using V2.5 chemistry (75 cycles, single-end 75 bp) or on the Illumina NovaSeq 6000 using the SP Reagent Kit (100 cycles, single-end 100bp) at the Genomics Core Facility at NYU. Control (without synthetic Hox constructs) RNA-seq datasets were previously published: 0h in (*59*); and 48h/96h after RA/SAG in (*75*).

#### Hi-C

Cells were collected at 0h and 48h after RA treatment. Cells were divided into ∼1×10^6^ aliquots and crosslinked in a final concentration of 2% FA (vol/vol) in 1 × PBS for 10 min at room temperature. After quenching with Glycine, cells incubated on ice for 15 min, pelleted by centrifugation at 500 g, and frozen at -80°C. Hi-C was performed using the Arima-HiC workflow (Arima Genomics, San Diego, CA) by NYU Langone’s Genome Technology Center (RRID: SCR_017929).

### Sequencing data analysis

#### Custom references

Two modified versions of the mm10 genome and corresponding genome annotations, were created using the reform tool (https://reform.bio.nyu.edu/). The genome mm10_synHoxA was created by replacing mm10 chrX:52963048-52997452 (Hprt) with the sequence corresponding to 170kb *Enhancers* + *SynHoxA* delivered to the ICE landing pad. A second genome mm10_synHoxA_delHoxA was then created by removing the endogenous *HoxA* sequence that is deleted in the *HoxA* ^-/-^ mESC line (mm10 chr6:52151380-52285416).

The custom genome sequences, annotations and bowtie2 references can be found at: https://genome.med.nyu.edu/public/boekelab/SynHox_genomes/

#### Parental mESCs WGS analysis

WGS data were analyzed as previously described (*52*). Reads were demultiplexed with Illumina bcl2fastq v2.20 requiring a perfect match to indexing BC sequences. Illumina sequencing adapters were trimmed with Trimmomatic v0.39 (*76*). Reads were aligned to mm10 using BWA v0.7.17 (*77*). PCR duplicates were marked using samblaster v0.1.24 (*78*). Generation of per base coverage depth tracks and quantification was performed using BEDOPS v2.4.35 (*79*). Data were visualized using the University of California, Santa Cruz Genome Browser at coordinates mm10 chr6:52001889-52321810.

#### Assemblon BAC sequencing analysis

Reads were demultiplexed with Illumina bcl2fastq v2.20 requiring a perfect match to indexing BC sequences. Illumina sequencing adapters were trimmed with Trimmomatic v0.39. Reads were then mapped to mm10_synHoxA_delHoxA using bowtie2 v2.2.9 unless specified otherwise (*80*). Samtools was used to sort bam files and coverage tracks were generated using deeptools bamCoverage v3.2.1 (*81*) with bin size set to 1, ignoring duplicates (*81, 82*). Coverage was visualized in the Integrated Genomics Viewer (IGV) at coordinates chrX:52967270-53136430 (*83*).

Variants in *SynHoxA* assemblons were called relative to the rn6 rat reference genome. Sequencing data from assemblon BACs were mapped using the same pipeline described for parental mESCs (*52*). A modified rn6 genome was used as reference. Two sequences were masked: 1) the sequence corresponding to the mistaken duplication at *HoxA* (rn6 chr4: 82229539-82315425) and 2) an unplaced contig (4_KL567939v1_random) containing *HoxA* sequence. Variants were then called using a standard pipeline based on bcftools v1.9:

bcftools mpileup–redo-BAQ–adjust-MQ 50–gap-frac 0.05–max-depth 10000–max- idepth 200000 -a DP,AD–output-type u |

bcftools call–keep-alts –ploidy 1–multiallelic-caller -f GQ–output-type u

Raw pileups were filtered using:

bcftools norm–check-ref w–output-type u |

bcftools filter -i “INFO/DP>=10 & QUAL>=10 & GQ>=99 & FORMAT/DP>=10”–SnpGap 3–IndelGap 10–set-GTs .–output-type u |

bcftools view -i ’GT=”alt”’–trim-alt-alleles–output-type z

#### Capture-sequencing coverage analysis

All capture-seq data coming from assemblon delivery to WT mESCs was mapped to mm10_synHoxA whereas all data from *HoxA* ^-/-^ mESCs was mapped to mm10_synHoxA_delHoxA for coverage analysis. Reads were demultiplexed with Illumina bcl2fastq v2.20 requiring a perfect match to indexing BC sequences. Illumina sequencing adapters were trimmed with Trimmomatic v0.39. Reads were mapped using bowtie2 v2.2.9. Samtools was used to sort bam files and coverage tracks were generated using deeptools (*81*) bamCoverage v3.2.1 with bin size set to 1, ignoring duplicates. Coverage was visualized in the Integrated Genomics Viewer (IGV) coordinates chrX:52967270-53136430 (*83*).

#### Capture-sequencing integration site analysis

Bamintersect analysis was performed as previously described (*52*) with a few modifications. Briefly, capture-seq data were mapped to mm10 and independently to references containing the ICE landing pad sequence and the delivered assemblon sequence. Bamintersect identifies junctions by looking for read pairs where each read is mapped to a different reference.

To exclude spurious hits, certain sequences present in multiple contexts are masked. For example, the mouse Pgk1 promoter is found in the rtTA cassette integrated at Rosa26, at its endogenous location on chrX and at the 3’ end of the delivered assemblon driving G418 (Neo) resistance. Similarly, the SV40 pA signal is present downstream of both the GFP in the assemblon and G418^R^ (Neo) gene in the landing pad. Coordinates of masked sequences are: 1) Pgk1 promoter found in the rtTA cassette integrated at Rosa26 (mm10 chr6:113071694-113077114) 2) Pgk1 promoter – endogenous location (mm10 chrX:106186732-106187231) and 3) region of the mouse genome immediately downstream of the G418^R^ SV40 pA (chrX:52962425-52963047). In addition, the endogenous mouse *HoxA* sequence (mm10 chr6:52121869-52285368) is masked to eliminate hits that arise from cross mapping of highly conserved regions between the rat derived assemblon and mouse *HoxA*.

Reads with the same strand and mapping to within 500 bp of each other were clustered for reporting. Regions below 150 bp or with fewer than 2 reads/10M reads sequenced were excluded.

#### ChIP-seq data analysis

Fastq files were aligned to the mm10_synHoxA_delHoxA custom genome using Bowtie 2 (*80*), using options -p 20. Samtools (*82*) was used to create sorted bam files for inputs into the bamCoverage tool from deeptools (*81*) to create bigWig files, using options: -- binSize 1 --scaleFactor 0.001 --normalizeUsingRPKM --ignoreForNormalization chrX -p max --extendReads 100. The bigWig files were visualized using IGV (*83*).

To generate sliding window plots of ChIP-seq signal, bedtools makewindows was used to generate bins of 3kb sliding 300 bp. Bedtools coverage was then used to compute the mean coverage in each bin from the sorted bam files (*84*). Coverage was normalized across samples using RPKM that was calculated as: reads-per-bin/(number of mapped reads (in millions) * bin length (kb)). Mean value and standard deviation of replicates was then plotted using Python matplotlib.

Of note, the vector backbone for all integrated constructs was covered in H2K27me3 at all time points. This corroborates previous reports of bacterial sequence silencing in mammalian cells (Supplementary Figure 13) (*85*).

#### RNA-seq data analysis

Fastq files were aligned to the genome (mm10_synHoxA or mm10_synHoxA_delHoxA custom genomes) using HISAT2 (*86, 87*), using options: -p 20 -q --rna-strandness F. Mapped reads were assigned to annotated genes using the featureCount function in Rsubread (*88*), using options -s 2. Read counts were normalized using the ‘rlog’ or regularized log transformation in DESeq2 (*89*) and used as inputs for the Principal Component Analysis (PCA). The log2 fold change (FC) and adjusted p-value in gene expression levels between 24h, 48h, and 96h vs. 0h was estimated using DESeq2 and plotted using ggplot (*90*). Normalized counts from DESeq2 were used to generate transgene/endogenous ratio plots, with the counts from endogenous locus being reduced by half to normalize the copy number between HoxA clusters. RNAseq track visualization in Suppl. Fig. 6 was performed using combine tracks tool in IGV (*83*) on bigwigs generated with deeptools bamCoverage (*81*) and subsequent editing in Adobe Illustrator.

#### Hi-C data analysis

Hi-C data was aligned against the custom mm10_synHoxA_delHoxA genome using BWA mem (version 0.7.17) using parameters -M -t 4 and aligning each mate pair independently (*77*). Samtools (version 1.11) was used to sort mapped reads by read name, and the pair_reads.py script in mHiC was used to join mate-pairs into a paired-end SAM file (*82, 91*). Paired-end read counts were then binned at 10kbp resolution to create the genome-wide contact matrix. The TAD calls displayed in Supplementary Figure 8 were produced using HiCseg (version 1.1) (*92*). The HiCseg_linkC_R function in HiCseg was provided with the segment of the 10kbp-resolution contact matrix corresponding to coordinates chrX:43000000-63000000 (mm10_synHoxA_delHoxA custom genome), and the following parameters: nb_change_max=100, distrib=“G”, model=“D”. The heatmap was then plotted at coordinates chrX:52400000-53600000 and chrX:52800000-53300000, with maximum color intensity set at a contact frequency of 40.

---

**SuppFigure 1:**
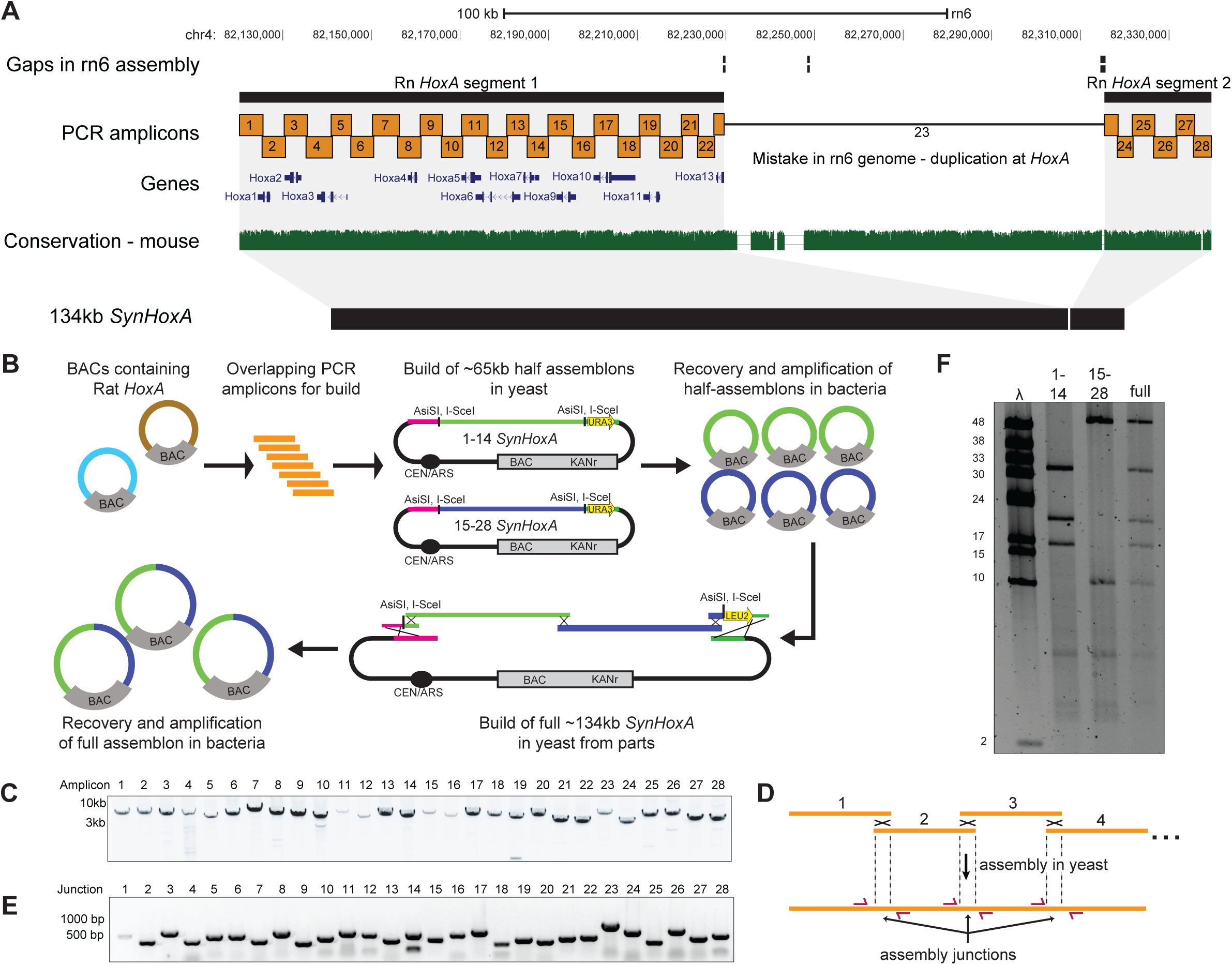
Build of 134kb *SynHoxA* assemblon. (A) Layout of rat *HoxA* locus in the rn6 genome assembly. The rn6 genome includes an erroneous duplication at the *HoxA* locus between gaps in the assembly. The *SynHoxA* assemblon sequence is based on bringing together the two ‘separate’ RnHoxA segments. The sequence was segmented into 28 ∼5kb PCR amplicons with terminal homology of ∼200bp to adjacent amplicons. Conservation to the mouse genome is depicted using the multiz track from the UCSC genome browser. (B) Schematic depicting the assembly workflow for the 134kb *SynHoxA* assemblon. BACs containing Rat *HoxA* were used as PCR template to generate 28 segments tiling the entire *HoxA* locus. These segments were co-transformed into yeast with appropriate linkers and assembly vector to build two ∼65kb half assemblons into centromeric yeast-bacteria shuttle vectors. These half assemblons are recovered to bacteria and amplified. Full 134kb assemblon was built from half assemblons after releasing them from the vector using terminal restriction enzymes (*AsiSI*) and transforming into yeast. Full assemblon was then recovered from yeast into bacteria for amplification and verification. (C) Agarose gel of the 28 PCR amplicons that tile the 134kb *SynHoxA* assemblon. (D) Strategy to PCR-screen yeast colonies derived from assembly experiments. Primers (red arrows) span assembly junctions and test presence/absence of amplicons in many yeast colonies. Reproduced from ref (47) with permission from authors. (E) Agarose gel showing one yeast colony carrying the full 134kb *SynHoxA* assemblon verified manually for the presence of all assembly junctions, using the strategy outlined in panel D. (F) Half and Full 134kb *SynHoxA* assemblon BACs purified from *E.coli* were digested with PvuI and separated using field inversion gel electrophoresis (FIGE). Lambda monocut ladder sizes are indicated in kb. Band sizes correspond to expected fragments.

**SuppFigure 2:**
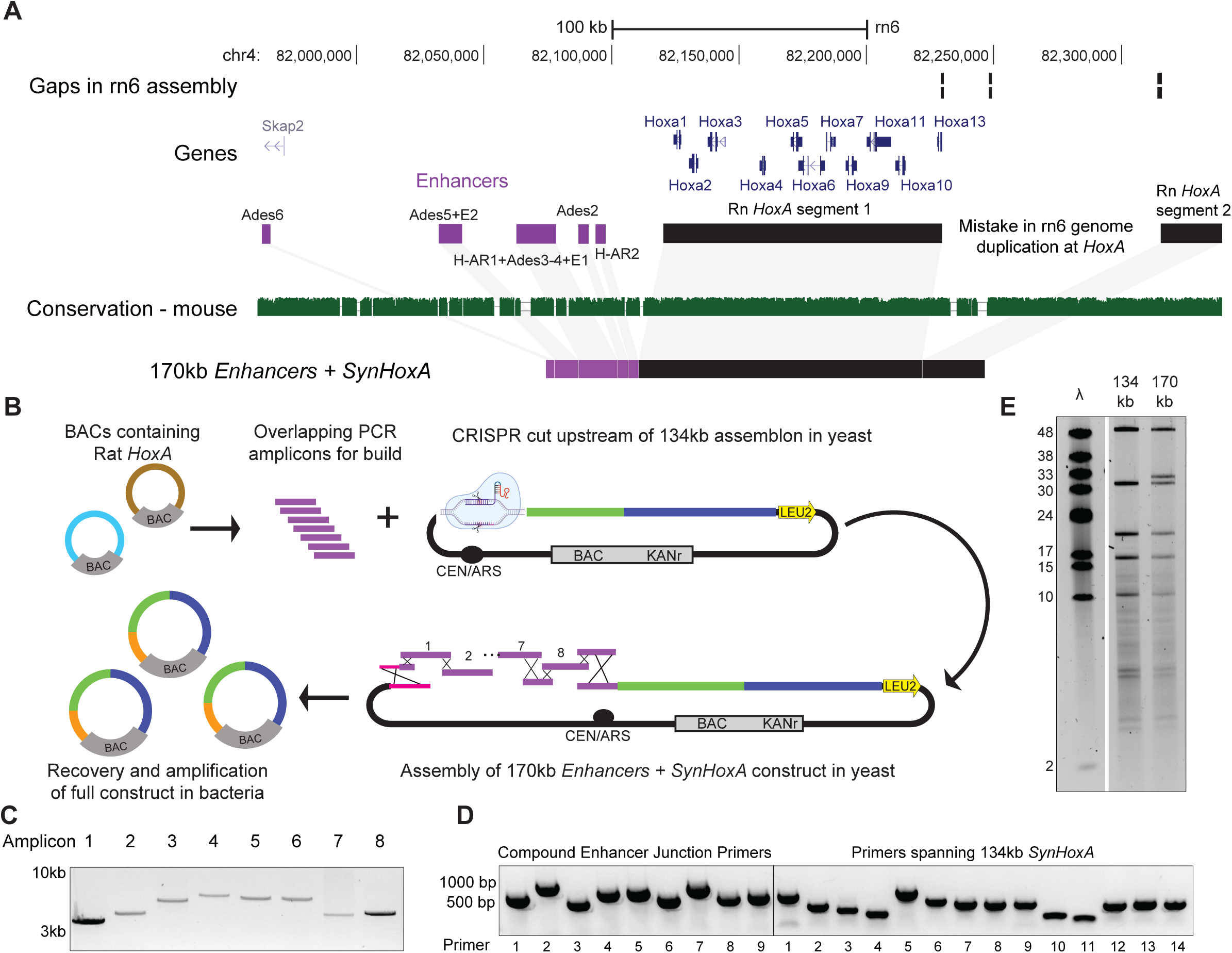
Build of 170kb *Enhancers+SynHoxA* assemblon. (A) Layout of rat *HoxA* locus from the rn6 genome assembly depicting genes, Rn *HoxA* cluster segments in black and previously identified distal enhancers in purple. The *Enhancers+SynHoxA* assemblon sequence is made by stringing all the enhancers directly upstream of the *SynHoxA* assemblon sequence. Conservation to mouse genome is depicted using multiz track from the UCSC genome browser. (B) PCR amplicons tiling enhancer sequences were generated from Rat *HoxA* BACs and co-transformed into a yeast strain containing the 134kb *SynHoxA* assemblon with a gRNA vector targeting the left terminus of the 134kb assemblon. The enhancer PCR amplicons were used to repair this break, resulting in the construction of the 170kb *Enhancers+SynHoxA* assemblon. Assemblon was recovered into bacteria for amplification and verification. (C) Agarose gel of the 8 PCR amplicons containing enhancer sequences. (D) Agarose gel showing one yeast colony tested for the presence of novel enhancer assembly junctions and with primers spanning 134kb *SynHoxA*. (E) 134kb and 170kb assemblon BACs purified from *E.coli* were digested with *PvuI* and separated using FIGE. Lambda monocut ladder sizes are indicated in kb. Band sizes correspond to expected fragments.

**SuppFigure 3:**
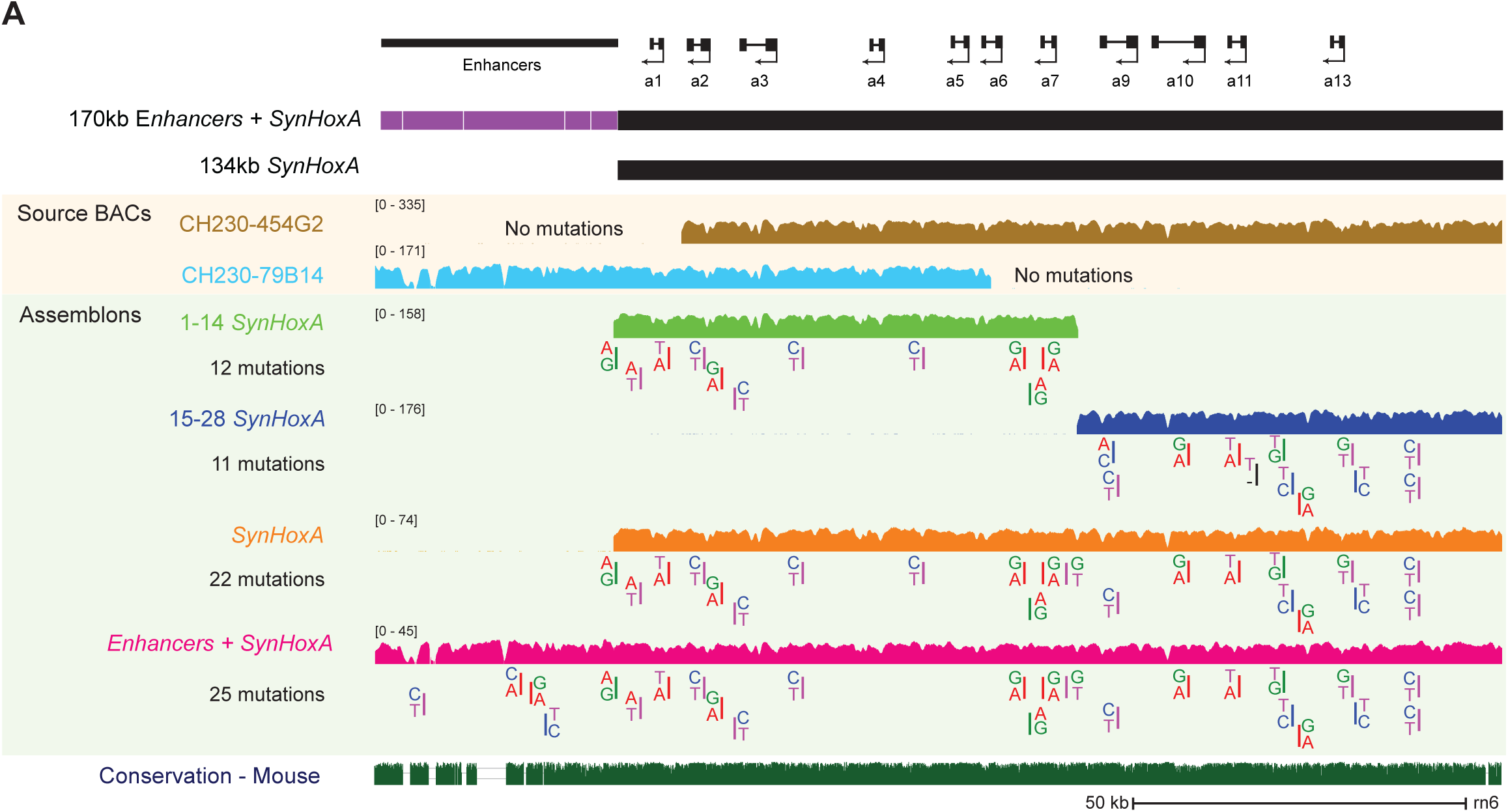
Mutations in *SynHoxA* assemblon that are generated during construction. Sequencing data from various stages of assemblon construction: source rat BACs, half-assemblon BACs and full-assemblon BACs. Below each coverage track, variant positions in comparison to the reference rn6 genome are depicted with reference allele (top) and variant allele (bottom). Data shown here are aligned to the rat reference genome rn6 and smoothed over 500bp.

**SuppFigure 4:**
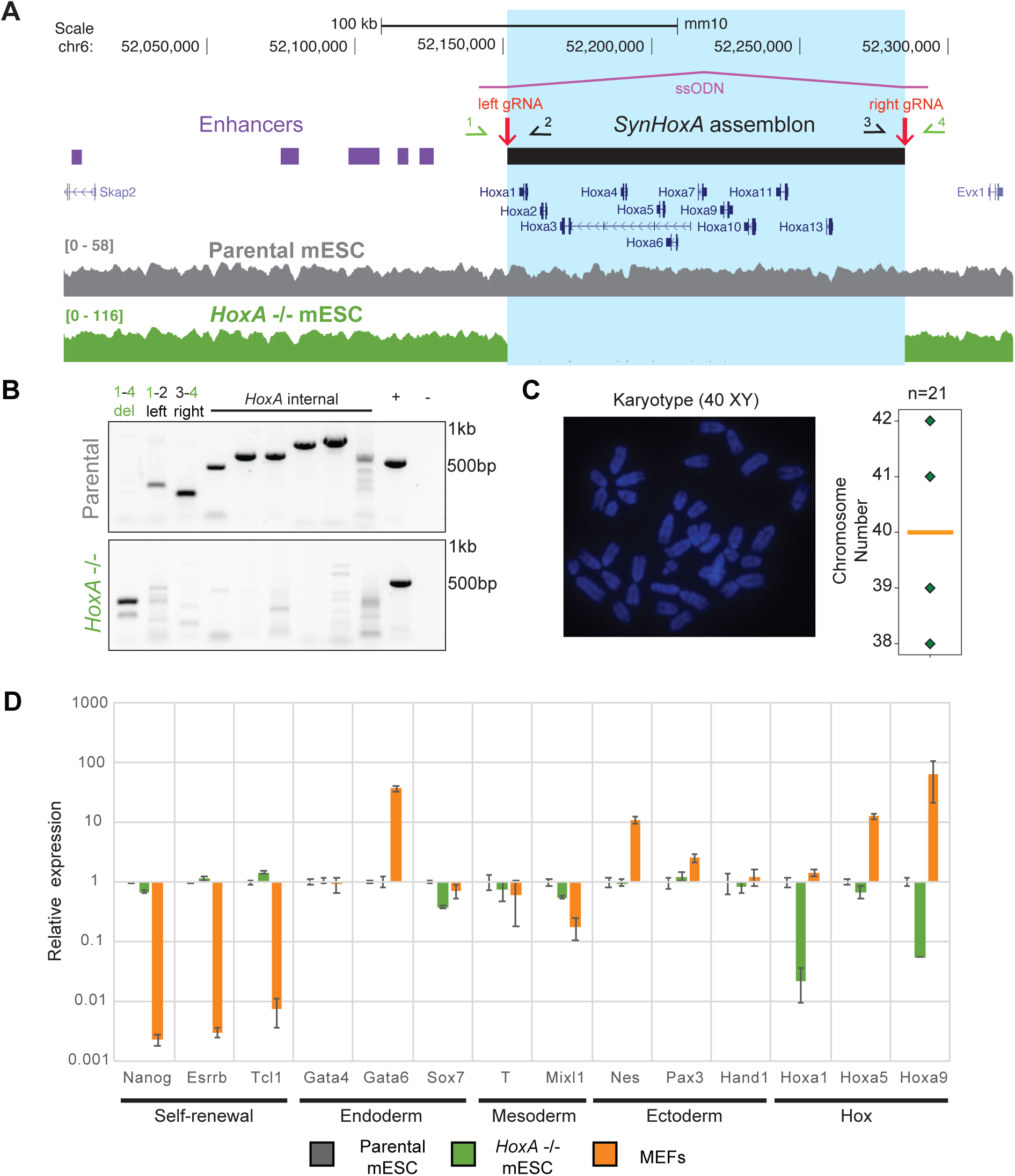
Generation of *HoxA* ^-/-^ mESCs by CRISPR/Cas9 induced deletion. (A) Top, layout of mouse *HoxA* locus with sequence corresponding to the 134kb *SynHoxA* assemblon marked with a black box and enhancers depicted with purple boxes. Deletion was induced by targeting with two guide RNAs (Left and Right) and by providing a single stranded oligo donor (ssODN). Primers used for genotyping PCR are depicted (1–4). Bottom, whole genome sequencing data for parental and *HoxA* ^-/-^ mESCs aligned to mouse reference genome mm10 and smoothed over 2000bp. (B) Genotyping PCR for verifying *HoxA* ^-/-^ mESCs in comparison to parental cells using deletion specific primer pairs depicted in (A) and primers internal to the deletion. +, positive control with primers amplifying mouse *Furin* locus; –, no primer control. (C) Karyotyping of *HoxA* ^-/-^ mESCs by metaphase spreads to confirm euploidy. One representative spread is shown. Quantification of 21 spreads is presented on the right as a boxplot, confirming that the cells have an euploid mean chromosome number of 40. (D) qRT-PCR data from parental mESCs, *HoxA ^-/-^* mESCs and Mouse Embryonic Fibroblasts (MEFs) for a range of genes is shown. Relative expression is presented as normalized to *Gapdh* and parental mESCs.

**SuppFigure 5:**
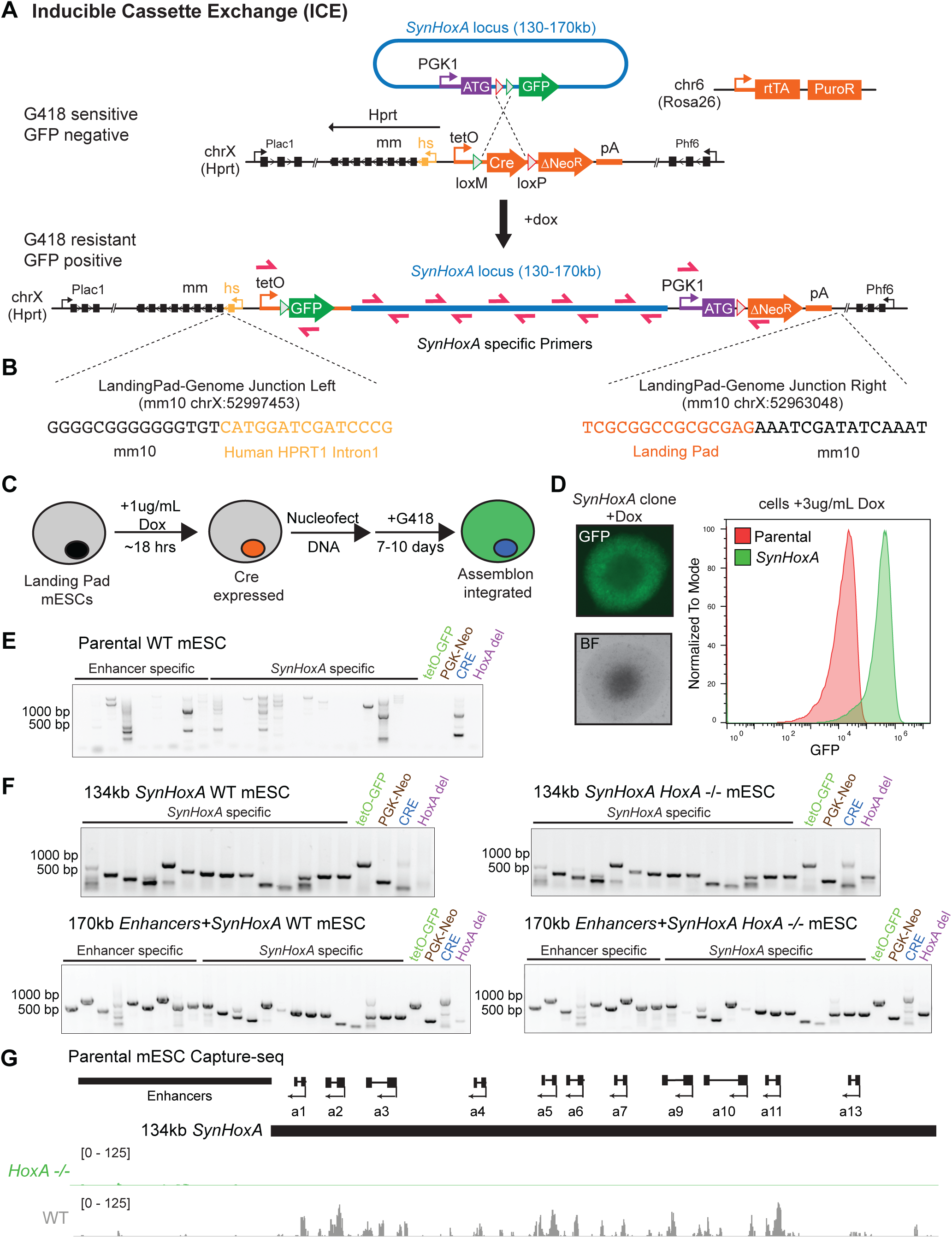
Delivery and PCR genotyping of *SynHoxA* variant assemblons. (A) Schematic of Inducible Cassette Exchange (ICE) for site-specific delivery of assemblons to mESCs. A resident landing pad, integrated at the mouse *Hprt* locus, contains a *Cre* recombinase gene, driven by a tetracycline inducible promoter (TRE) and is flanked by heterotypic loxM and loxP sites. A promoter-less Neomycin resistance gene lacking a start codon (ΔNeo) is found downstream of the *Cre*. The reverse tetracycline transactivator (rtTA) is expressed from the *Rosa26* locus. The assemblon vector contains a delivery cassette (PGK1-ATG-loxP-loxM-GFP). During cassette exchange, two Cre mediated recombination events result in the placement of GFP under the control of the tetO promoter, donation of the *PGK1* promoter and ATG start codon to ΔNeo, as well as loss of the *Cre* gene. This gives rise to G418-resistant (the Neo gene confers G418 resistance), GFP positive cells. Primers used for genotyping are indicated in red. (B) Sequence and coordinates of landing pad junctions with the mouse genome. (C) mESCs bearing the ICE landing pad on the X chromosome are treated with 1 µg/mL Doxycycline to induce *Cre* expression. DNA is nucleofected and cells are selected with G418 for 7-10 days until clones are grown out. Clones are then picked for genotyping and sequencing. (D) Left, an image of a representative *SynHoxA* clone is shown. Right, flow cytometry data from parental cells (red) and *SynHoxA* cells (green) after treatment with 3 µg/mL doxycycline. (E-F) Agarose gels showing genotyping of parental WT mESCs (E) and from clones arising from delivery of 134kb *SynHoxA* and 170kb *Enhancers+SynHoxA* to WT and *HoxA* ^-/-^ mESCs (F). Clones were screened using *SynHoxA*-specific primers that span the length of the assembly, primers that span novel junctions formed with the genome (tetO-GFP, PGK-Neo) and primers that confirm overwriting of the *Cre* gene. In addition, the presence or absence of the endogenous *HoxA* cluster deletion was confirmed by deletion-specific primers (see Supplementary Figure 4). (G) Capture-sequencing data generated from parental WT and *HoxA* ^-/-^ mESCs using rat *HoxA* sequence as bait. Data are aligned to custom mouse reference genomes and normalized for sequencing depth. There is minimal cross-mapping between endogenous *HoxA* and *SynHoxA* loci.

**SuppFigure 6:**
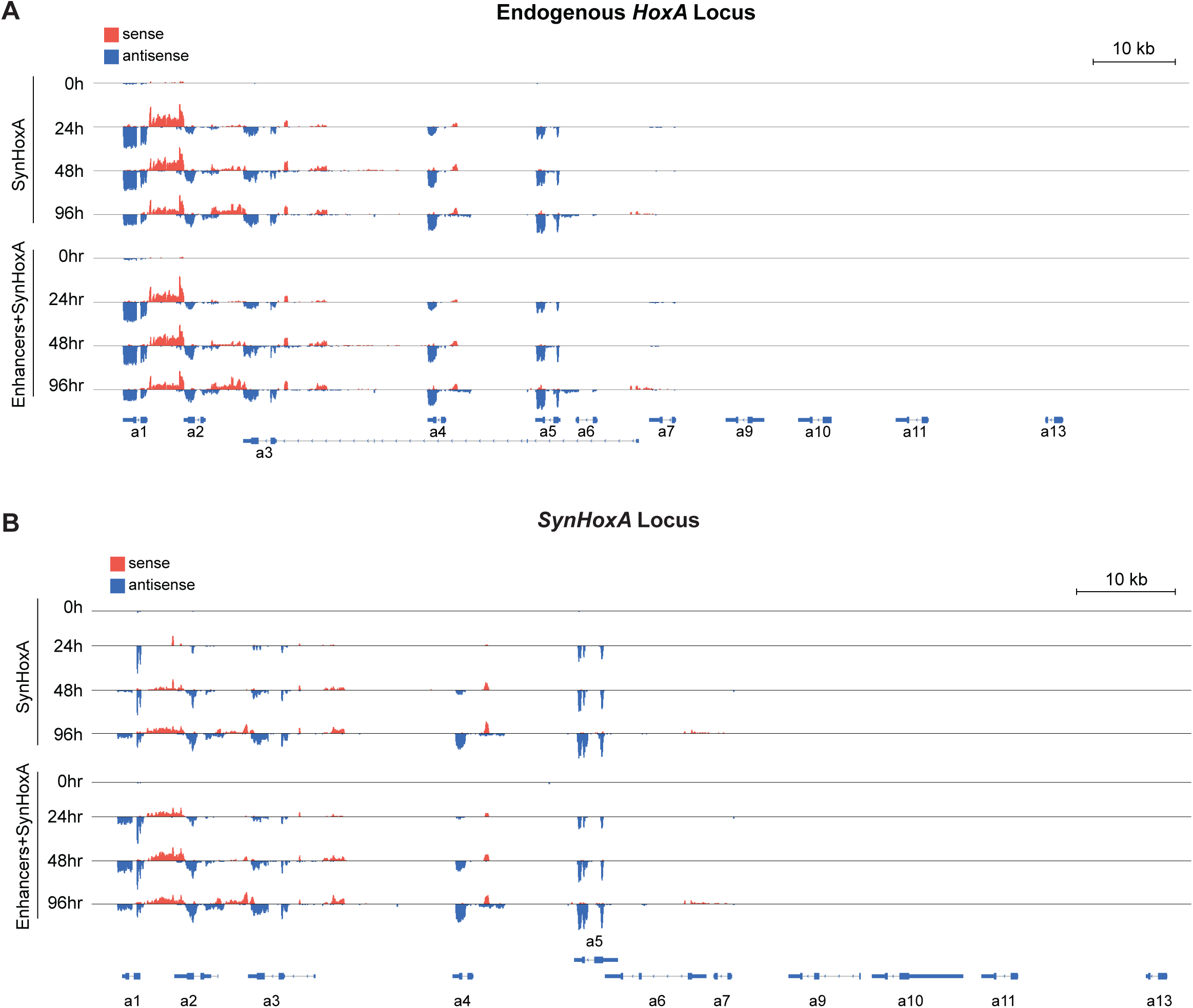
Visualization of raw gene expression data through differentiation. RNA-seq data through RA induced differentiation of *SynHoxA* and Enhancers + *SynHoxA* lines with an intact endogenous *HoxA* cluster are presented. Reads mapping to the endogenous cluster are shown in (A) and reads mapping to the *SynHoxA* cluster are in (B). Reads mapping to the sense strand are in red and antisense reads are in blue. Expression of both coding and non-coding transcripts was observed.

**SuppFigure 7:**
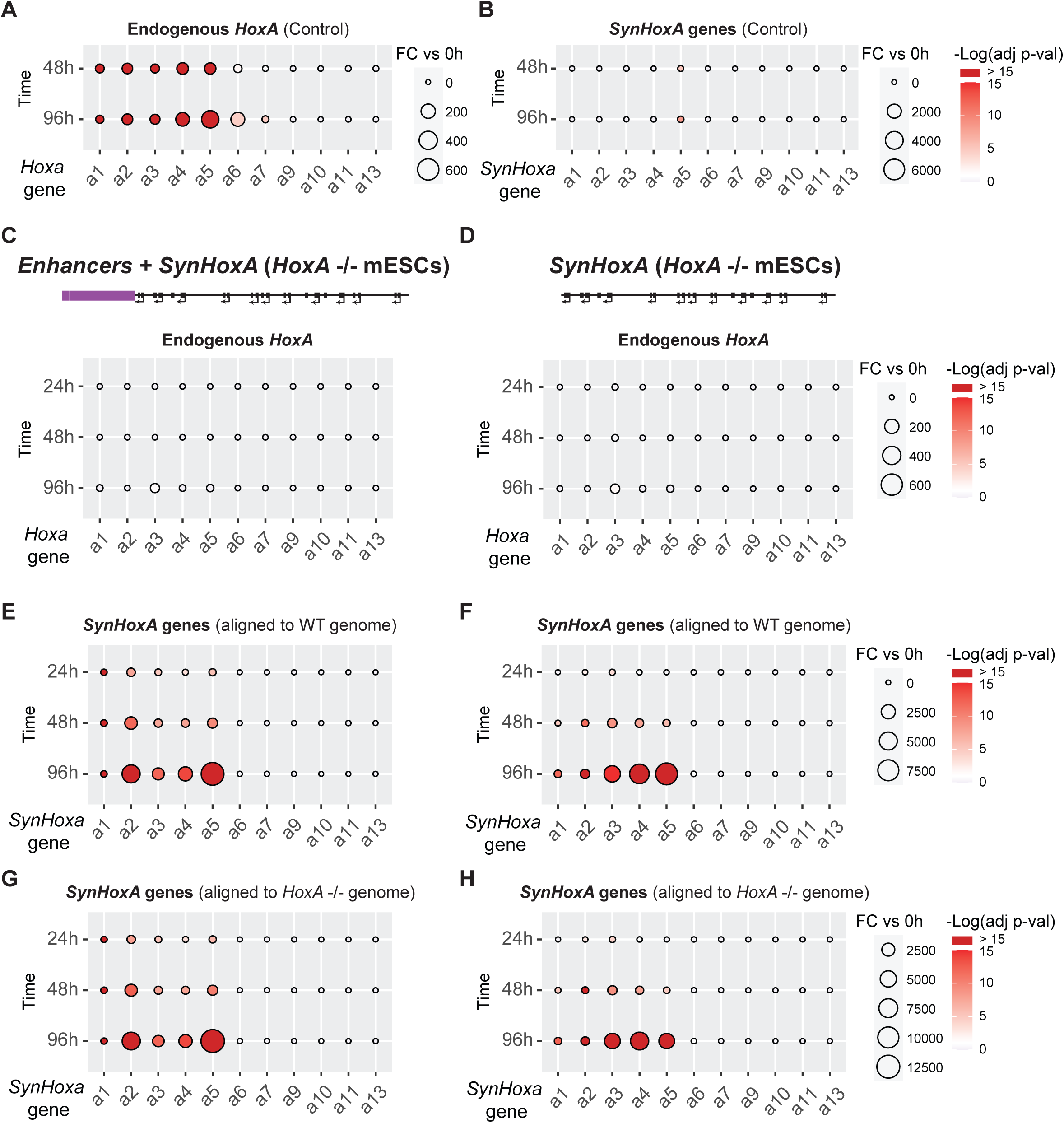
No mapping issues are revealed in the RNA-seq analysis. (A-B) The fold change of *SynHoxA* or endogenous mouse *HoxA* genes from RNA-seq data during differentiation in control lines, which do not contain *SynHoxA* variant clusters. The endogenous *HoxA* cluster induces the correct set of genes, and has some weak *Hoxa6* expression at the latest time point. Only a minor amount of reads map to *SynHoxa5* at 96h in control lines with no *SynHoxA* variant clusters. RNA-seq data are aligned to a modified mm10 mouse genome which contains the *SynHoxA* sequence inserted at the *Hprt* locus. Only uniquely mapped reads were kept and used in downstream analysis. (C-H) mapping tests similar to that in (A) for lines lacking the endogenous *HoxA* cluster but containing Enhancers + *SynHoxA* (C, E, G) and *SynHoxA* (D, F, H) at *Hprt.* (C-D) No mapping was observed to endogenous *HoxA* genes in cell lines lacking the endogenous *HoxA* cluster as expected. (E-H) Mapping RNA-seq data to a genome either containing (WT, E-F) or lacking the endogenous *HoxA* cluster (*HoxA ^-/-^*, G-H) did not affect the analysis.

**SuppFigure 8:**
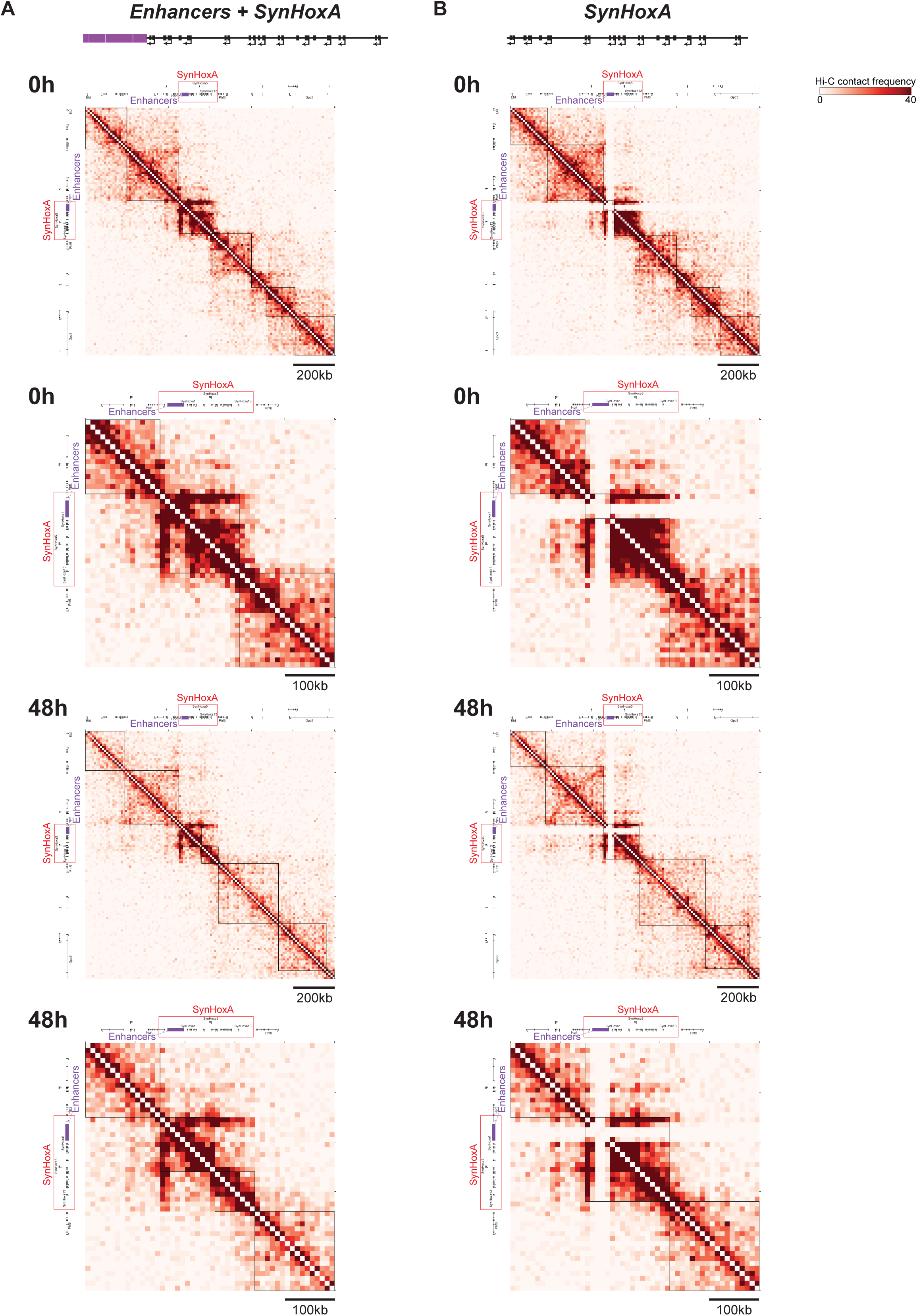
Ectopic *SynHoxA* clusters self-organize in 3D during differentiation. Heatmaps of Hi-C data during MN differentiation from mESCs lacking the endogenous *HoxA* cluster that contain either *Enhancers + SynHoxA* (A) or *SynHoxA* (B) at *Hprt*. Black lines indicate topological boundaries called by an unbiased algorithm, HiSeg2.0. A topological boundary formed between *SynHoxa5* and *SynHoxa6* in *Enhancers + SynHoxA*, mirroring endogenous organization.

**SuppFigure 9:**
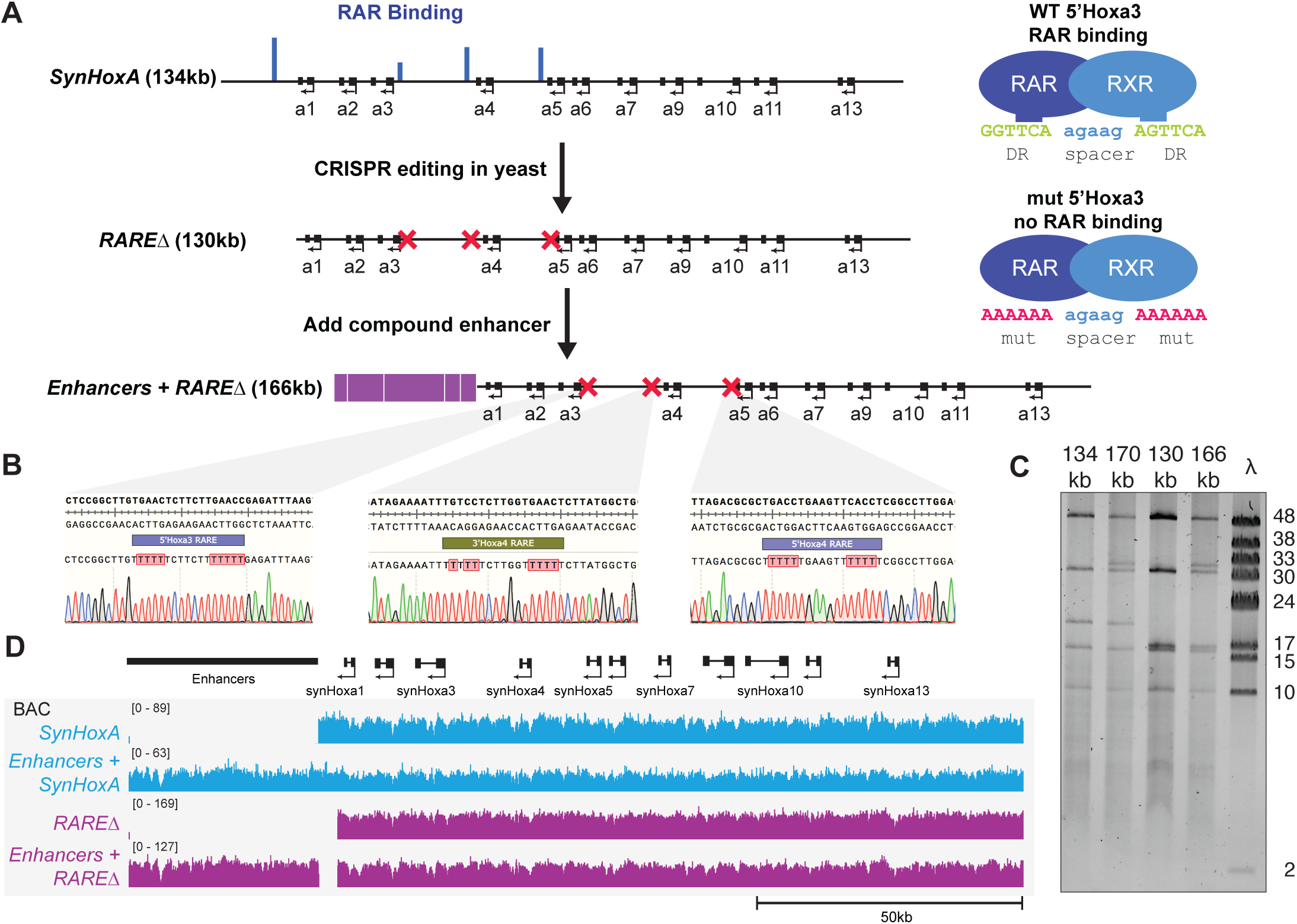
Build of *RAREΔ SynHoxA* assemblons. (A) Schematic of assembly strategy for 130kb *RAREΔ SynHoxA* and 166kb Enhancers + *RAREΔ SynHoxA*. Nature of the RARE mutations is shown on the right. RAR binding data comes from previously published reports. (see Methods) (B) Sanger sequencing traces confirmed precise CRISPR editing of RAREs in yeast. (C) *SynHoxA* assemblon BACs purified from *E.coli* were digested with *PvuI* and separated using FIGE. Lambda monocut ladder sizes are indicated in kb. Bands correspond to expected fragment lengths. (D) Sequencing data of assemblon BACs purified from *E. coli* aligned to a custom mm10 reference genome. Positions of the enhancers and protein coding genes are shown in black.

**SuppFigure 10:**
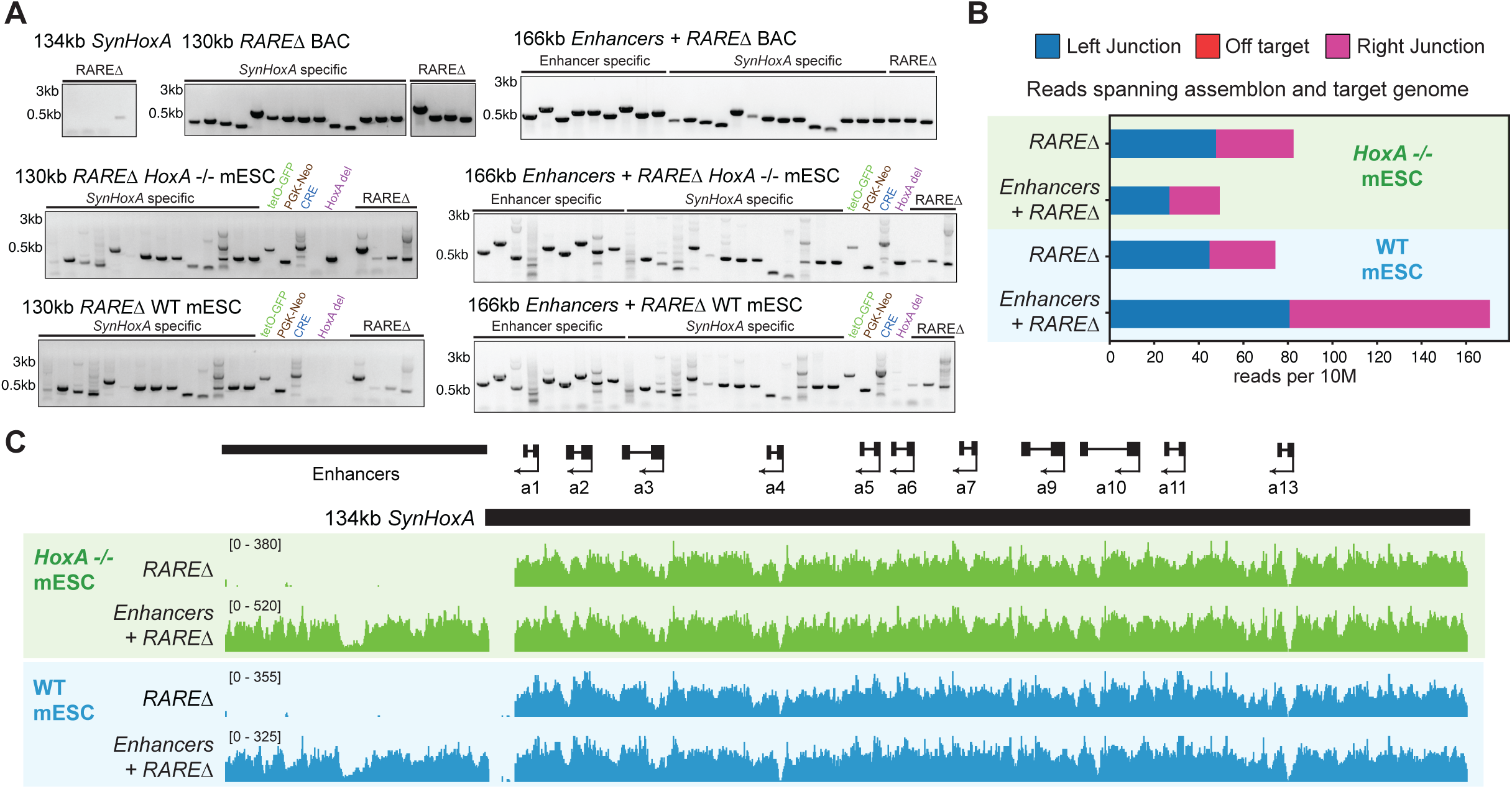
Delivery of *RAREΔ* assemblons to WT and *HoxA* ^-/-^ mESCs. (A) Genotyping PCRs separated on an agarose gel. RAREΔ specific primers are used to verify presence of the RARE mutations. Clones were also screened using *SynHoxA* specific primers that span the length of the assembly, primers specific to enhancer junctions, primers that span novel junctions formed with the genome (tetO-GFP, PGK-Neo) and primers that confirm overwriting of the *Cre* gene. In addition, the presence or absence of the endogenous *HoxA* cluster deletion was confirmed using deletion specific primers. (B) Only the expected junctions spanning the synthetic assemblon and the host genome were observed in next generation sequencing data with no off-target integrations. (C) Positions of the enhancers and protein coding genes are shown in black. Capture sequencing data is shown from WT and *HoxA* ^-/-^ mESC clones arising from delivery of *RAREΔ SynHoxA* assemblons. Sequencing data shown here are aligned to a custom reference genome (see Methods).

**SuppFigure 11:**
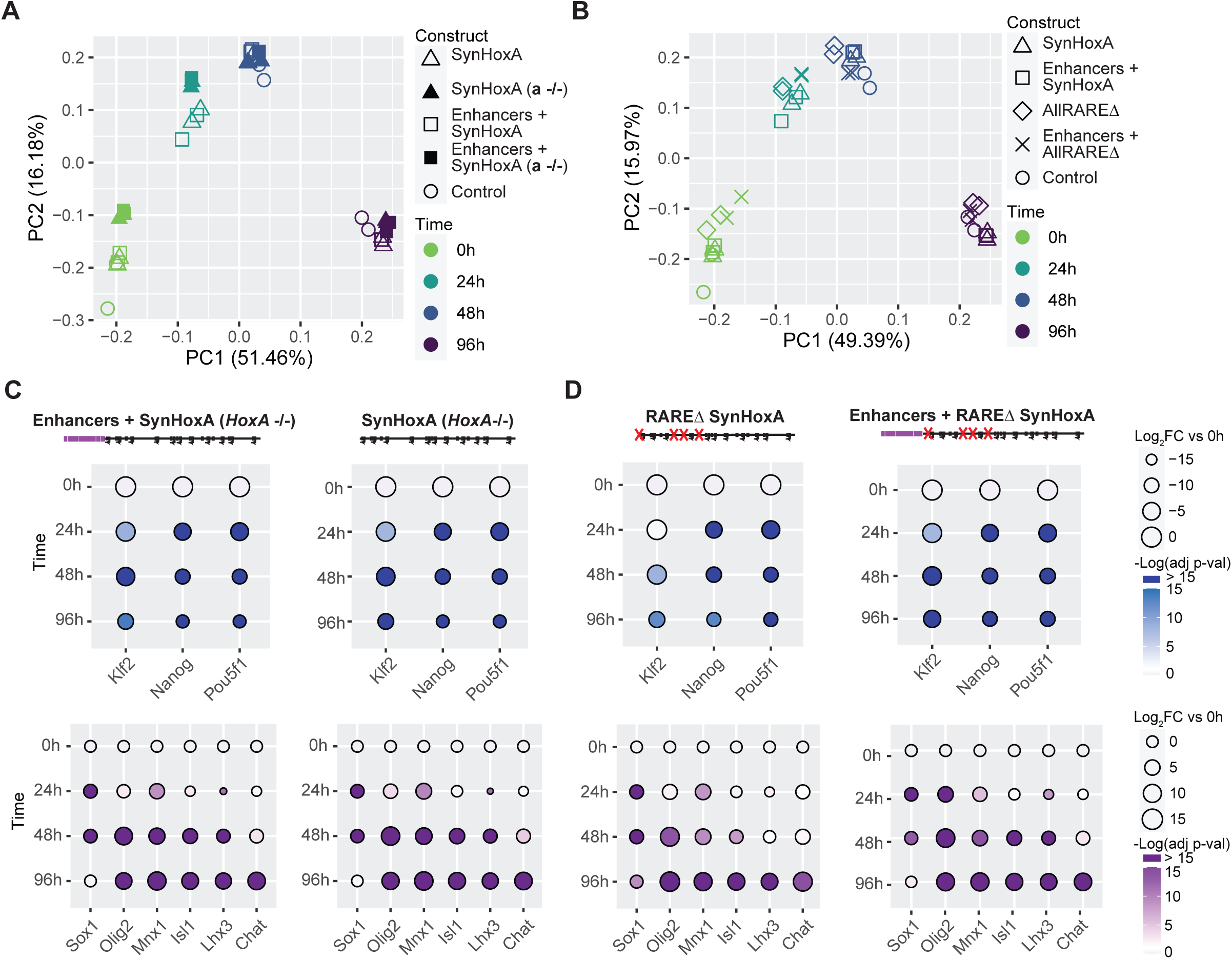
mESCs with *SynHoxA* assemblons differentiate well into MNs. (A-B) Principal Component Analysis (PCA) of the RNA-seq datasets reveals clustering largely by time during the differentiation protocol (each data point represents independent differentiations). This is true regardless of genetic background (A) and ectopic *SynHoxA* variants that are integrated (B). (C-D) The log2 fold change of pluripotency markers and MN differentiation genes from RNA-seq data (n=2). Pluripotency markers were downregulated and MN markers were upregulated during differentiation as expected for *SynHoxA* mESCs lacking the mouse *HoxA* cluster (C) and for *RAREΔ SynHoxA* mESCs (D).

**SuppFigure 12:**
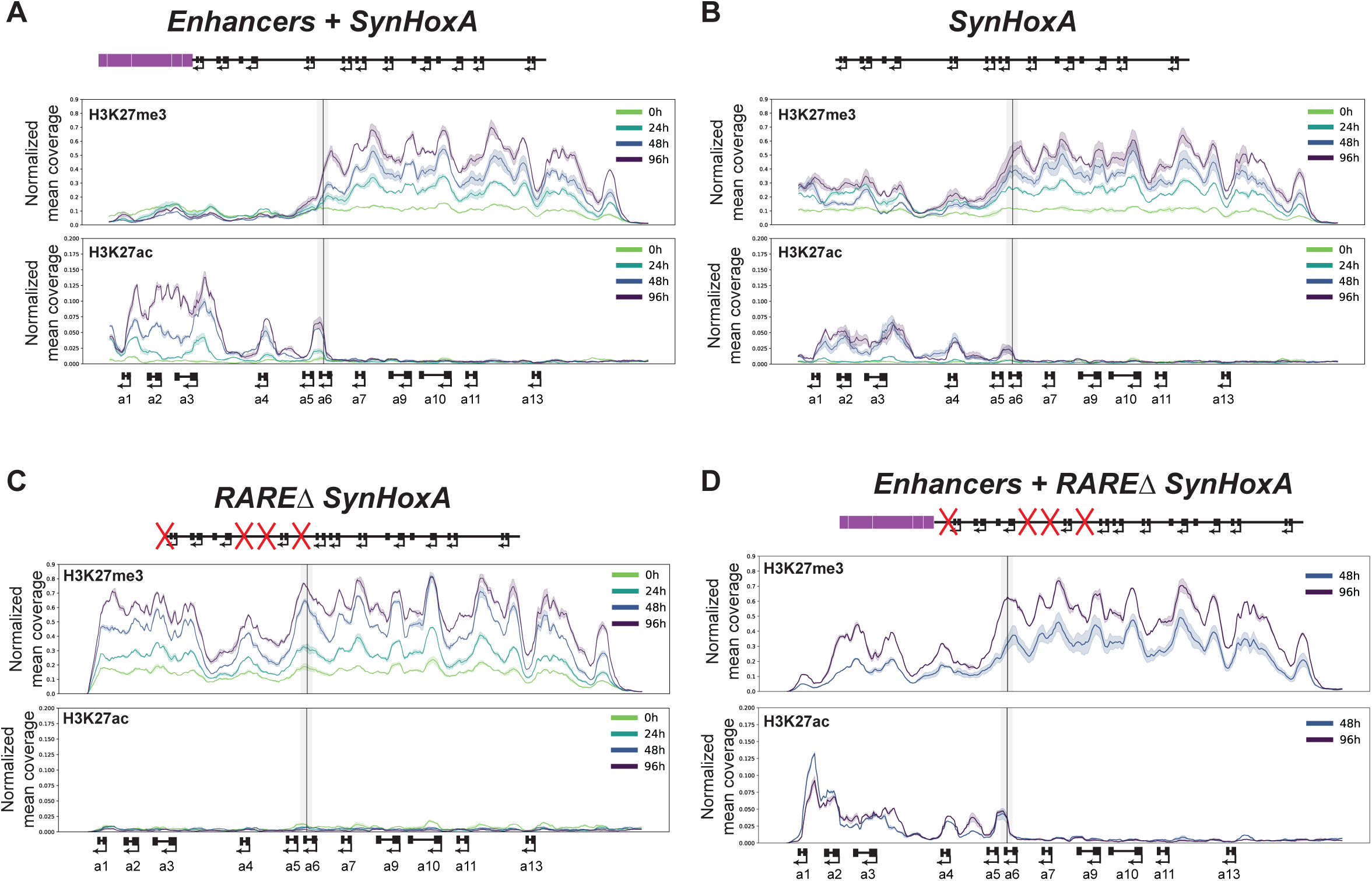
H3K27me3 and H3K27Ac distribution across *SynHoxA* during MN differentiation. RPKM normalized mean coverage on 3kb windows sliding 300bp across *SynHoxA* of H3K27me3 and H3K27Ac ChIP-seq data for each *SynHoxA* cell line. Data are aligned to a custom reference genome (see Methods).

**SuppFigure 13:**
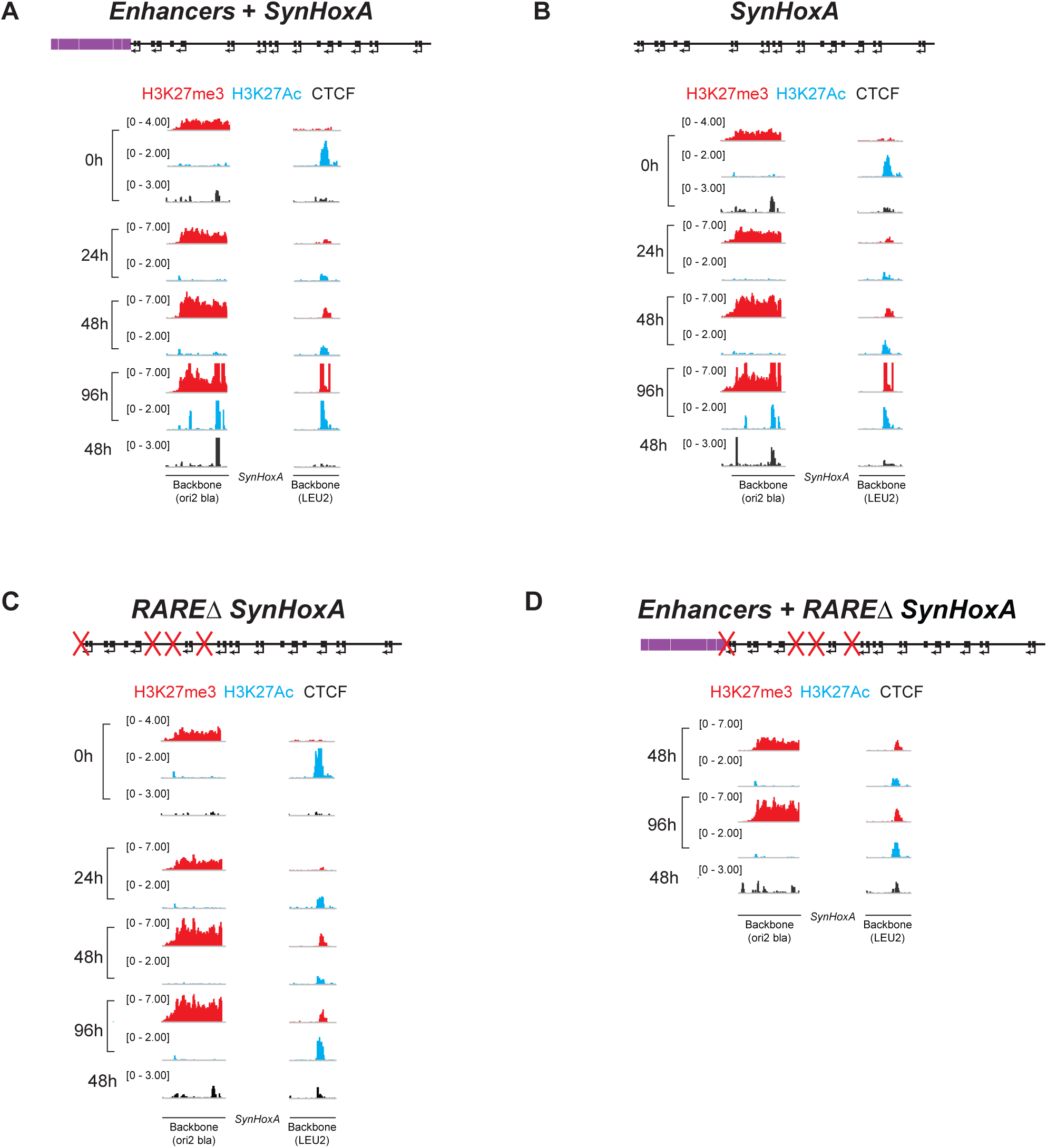
Vector backbones recruit chromatin modifications. ChIP-seq data mapping to the vector backbones associated with all *SynHoxA* clusters is presented at all time points analyzed. As part of the ICE delivery, the two parts of the vector backbone are separated by the *SynHoxA* cluster. Bacterial derived sequences were covered with the repressive chromatin modification H3K27me3 (red). The yeast *LEU2* gene recruited the activating chromatin modification H3K27Ac (blue) and H3K27me3. Some inconsistent CTCF (black) recruitment was observed. ChIP-seq data shown here are aligned to a custom mm10 reference genome (see Methods).

## REFERENCES

1. W. de Laat, D. Duboule, Topology of mammalian developmental enhancers and their regulatory landscapes. Nature 502, 499–506 (2013).

2. J. Deschamps, D. Duboule, Embryonic timing, axial stem cells, chromatin dynamics, and the Hox clock. Genes Dev 31, 1406–1416 (2017).

3. J. Deschamps, J. van Nes, Developmental regulation of the Hox genes during axial morphogenesis in the mouse. Development 132, 2931–2942 (2005).

4. M. Kmita, D. Duboule, Organizing axes in time and space; 25 years of colinear tinkering. Science 301, 331–333 (2003).

5. W. McGinnis, R. Krumlauf, Homeobox genes and axial patterning. Cell 68, 283–302 (1992).

6. E. B. Lewis, A gene complex controlling segmentation in Drosophila. Nature 276, 565–570 (1978).

7. D. Duboule, The rise and fall of Hox gene clusters. Development 134, 2549–2560 (2007).

8. D. Duboule, G. Morata, Colinearity and functional hierarchy among genes of the homeotic complexes. Trends Genet 10, 358–364 (1994).

9. T. Montavon, N. Soshnikova, Hox gene regulation and timing in embryogenesis. Semin Cell Dev Biol 34, 76–84 (2014).

10. T. Montavon, D. Duboule, Chromatin organization and global regulation of Hox gene clusters. Philos Trans R Soc Lond B Biol Sci 368, 20120367 (2013).

11. T. Montavon et al., A regulatory archipelago controls Hox genes transcription in digits. Cell 147, 1132–1145 (2011).

12. S. Berlivet et al., Clustering of tissue-specific sub-TADs accompanies the regulation of HoxA genes in developing limbs. PLoS Genet 9, e1004018 (2013).

13. N. Lonfat, T. Montavon, F. Darbellay, S. Gitto, D. Duboule, Convergent evolution of complex regulatory landscapes and pleiotropy at Hox loci. Science 346, 1004–1006 (2014).

14. K. Cao et al., SET1A/COMPASS and shadow enhancers in the regulation of homeotic gene expression. Genes Dev 31, 787–801 (2017).

15. B. De Kumar et al., Analysis of dynamic changes in retinoid-induced transcription and epigenetic profiles of murine Hox clusters in ES cells. Genome Res 25, 1229–1243 (2015).

16. R. Neijts et al., Polarized regulatory landscape and Wnt responsiveness underlie Hox activation in embryos. Genes Dev 30, 1937–1942 (2016).

17. N. Lin, J. Dang, T. M. Rana, Haunting the HOXA Locus: Two Faces of lncRNA Regulation. Cell Stem Cell 16, 449–450 (2015).

18. R. Margueron, D. Reinberg, The Polycomb complex PRC2 and its mark in life. Nature 469, 343–349 (2011).

19. E. O. Mazzoni et al., Saltatory remodeling of Hox chromatin in response to rostrocaudal patterning signals. Nat Neurosci 16, 1191–1198 (2013).

20. D. Noordermeer et al., The dynamic architecture of Hox gene clusters. Science 334, 222–225 (2011).

21. N. Soshnikova, D. Duboule, Epigenetic temporal control of mouse Hox genes in vivo. Science 324, 1320–1323 (2009).

22. C. Nolte, B. De Kumar, R. Krumlauf, Hox genes: Downstream “effectors” of retinoic acid signaling in vertebrate embryogenesis. Genesis, e23306 (2019).

23. S. Mahony et al., Ligand-dependent dynamics of retinoic acid receptor binding during early neurogenesis. Genome Biol 12, R2 (2011).

24. V. Narendra et al., CTCF establishes discrete functional chromatin domains at the Hox clusters during differentiation. Science 347, 1017–1021 (2015).

25. V. Narendra, M. Bulajic, J. Dekker, E. O. Mazzoni, D. Reinberg, CTCF-mediated topological boundaries during development foster appropriate gene regulation. Genes Dev 30, 2657–2662 (2016).

26. J. R. Dixon et al., Topological domains in mammalian genomes identified by analysis of chromatin interactions. Nature 485, 376–380 (2012).

27. N. Ostrov et al., Technological challenges and milestones for writing genomes. Science 366, 310–312 (2019).

28. M. Gasperini, L. Starita, J. Shendure, The power of multiplexed functional analysis of genetic variants. Nat Protoc 11, 1782–1787 (2016).

29. N. Heintz, BAC to the future: the use of bac transgenic mice for neuroscience research. Nat Rev Neurosci 2, 861–870 (2001).

30. K. R. Peterson et al., Transgenic mice containing a 248-kb yeast artificial chromosome carrying the human beta-globin locus display proper developmental control of human globin genes. Proc Natl Acad Sci U S A 90, 7593–7597 (1993).

31. K. R. Peterson et al., Use of yeast artificial chromosomes (YACs) in studies of mammalian development: production of beta-globin locus YAC mice carrying human globin developmental mutants. Proc Natl Acad Sci U S A 92, 5655–5659 (1995).

32. F. G. Liberante, T. Ellis, From kilobases to megabases: Design and delivery of large DNA constructs into mammalian genomes. Current Opinion in Systems Biology 25, 1–10 (2021).

33. H. A. Wallace et al., Manipulating the mouse genome to engineer precise functional syntenic replacements with human sequence. Cell 128, 197–209 (2007).

34. N. S. McCarty, A. E. Graham, L. Studena, R. Ledesma-Amaro, Multiplexed CRISPR technologies for gene editing and transcriptional regulation. Nat Commun 11, 1281 (2020).

35. K. Kraft et al., Deletions, Inversions, Duplications: Engineering of Structural Variants using CRISPR/Cas in Mice. Cell Rep 10, 833–839 (2015).

36. D. G. Gibson et al., Complete chemical synthesis, assembly, and cloning of a Mycoplasma genitalium genome. Science 319, 1215–1220 (2008).

37. C. A. Hutchison, 3rd et al., Design and synthesis of a minimal bacterial genome. Science 351, aad6253 (2016).

38. J. Fredens et al., Total synthesis of Escherichia coli with a recoded genome. Nature 569, 514–518 (2019).

39. J. S. Dymond et al., Synthetic chromosome arms function in yeast and generate phenotypic diversity by design. Nature 477, 471–476 (2011).

40. N. Annaluru et al., Total synthesis of a functional designer eukaryotic chromosome. Science 344, 55–58 (2014).

41. L. A. Mitchell et al., Synthesis, debugging, and effects of synthetic chromosome consolidation: synVI and beyond. Science 355, (2017).

42. Z. X. Xie et al., “Perfect” designer chromosome V and behavior of a ring derivative. Science 355, (2017).

43. Y. Shen et al., Deep functional analysis of synII, a 770-kilobase synthetic yeast chromosome. Science 355, (2017).

44. Y. Wu et al., Bug mapping and fitness testing of chemically synthesized chromosome X. Science 355, (2017).

45. W. Zhang et al., Engineering the ribosomal DNA in a megabase synthetic chromosome. Science 355, (2017).

46. S. M. Richardson et al., Design of a synthetic yeast genome. Science 355, 1040–1044 (2017).

47. L. A. Mitchell et al., De novo assembly and delivery to mouse cells of a 101 kb functional human gene. Genetics, (2021).

48. N. Agmon et al., Yeast Golden Gate (yGG) for the Efficient Assembly of S. cerevisiae Transcription Units. ACS Synth Biol 4, 853–859 (2015).

49. M. Iacovino et al., Inducible cassette exchange: a rapid and efficient system enabling conditional gene expression in embryonic stem and primary cells. Stem Cells 29, 1580–1588 (2011).

50. M. Jasin, M. E. Moynahan, C. Richardson, Targeted transgenesis. Proceedings of the National Academy of Sciences 93, 8804–8808 (1996).

51. M. Gasperini et al., CRISPR/Cas9-Mediated Scanning for Regulatory Elements Required for HPRT1 Expression via Thousands of Large, Programmed Genomic Deletions. Am J Hum Genet 101, 192–205 (2017).

52. R. Brosh et al., A versatile platform for locus-scale genome rewriting and verification. Proc Natl Acad Sci U S A 118, (2021).

53. H. Wichterle, I. Lieberam, J. A. Porter, T. M. Jessell, Directed differentiation of embryonic stem cells into motor neurons. Cell 110, 385–397 (2002).

54. B. N. Davis-Dusenbery, L. A. Williams, J. R. Klim, K. Eggan, How to make spinal motor neurons. Development 141, 491–501 (2014).

55. H. Wichterle, M. Peljto, Differentiation of mouse embryonic stem cells to spinal motor neurons. Curr Protoc Stem Cell Biol Chapter 1, Unit 1H 1 1-1H 1 9 (2008).

56. M. Peljto, J. S. Dasen, E. O. Mazzoni, T. M. Jessell, H. Wichterle, Functional diversity of ESC-derived motor neuron subtypes revealed through intraspinal transplantation. Cell Stem Cell 7, 355–366 (2010).

57. M. Peljto, H. Wichterle, Programming embryonic stem cells to neuronal subtypes. Curr Opin Neurobiol 21, 43–51 (2011).

58. H. Jung et al., Global control of motor neuron topography mediated by the repressive actions of a single hox gene. Neuron 67, 781–796 (2010).

59. B. Aydin et al., Proneural factors Ascl1 and Neurog2 contribute to neuronal subtype identities by establishing distinct chromatin landscapes. Nat Neurosci 22, 897–908 (2019).

60. M. Bulajic et al., Differential abilities to engage inaccessible chromatin diversify vertebrate Hox binding patterns. Development 147, (2020).

61. E. Rodriguez-Carballo et al., Chromatin topology and the timing of enhancer function at the HoxD locus. Proc Natl Acad Sci U S A 117, 31231–31241 (2020).

62. G. Su et al., CTCF-binding element regulates ESC differentiation via orchestrating long-range chromatin interaction between enhancers and HoxA. J Biol Chem, 100413 (2021).

63. A. R. Amandio, L. Lopez-Delisle, C. C. Bolt, B. Mascrez, D. Duboule, A complex regulatory landscape involved in the development of mammalian external genitals. Elife 9, (2020).

64. M. Fernandez-Guerrero et al., Mammalian-specific ectodermal enhancers control the expression of Hoxc genes in developing nails and hair follicles. Proc Natl Acad Sci U S A 117, 30509–30519 (2020).

65. E. Rodriguez-Carballo et al., The HoxD cluster is a dynamic and resilient TAD boundary controlling the segregation of antagonistic regulatory landscapes. Genes Dev 31, 2264–2281 (2017).

66. O. Oksuz et al., Capturing the Onset of PRC2-Mediated Repressive Domain Formation. Mol Cell 70, 1149–1162 e1145 (2018).

67. K. A. Ganzinger, P. Schwille, More from less - bottom-up reconstitution of cell biology. J Cell Sci 132, (2019).

68. A. P. Liu, D. A. Fletcher, Biology under construction: in vitro reconstitution of cellular function. Nat Rev Mol Cell Biol 10, 644–650 (2009).

69. R. D. Gietz, R. H. Schiestl, High-efficiency yeast transformation using the LiAc/SS carrier DNA/PEG method. Nat Protoc 2, 31–34 (2007).

70. N. Agmon et al., Phylogenetic debugging of a complete human biosynthetic pathway transplanted into yeast. Nucleic Acids Res 48, 486–499 (2020).

71. J. E. DiCarlo et al., Genome engineering in Saccharomyces cerevisiae using CRISPR-Cas systems. Nucleic Acids Res 41, 4336–4343 (2013).

72. J. Sambrook, D. W. Russell, Isolation of BAC DNA from Small-scale Cultures. CSH Protoc 2006, (2006).

73. A. Untergasser et al., Primer3--new capabilities and interfaces. Nucleic Acids Res 40, e115 (2012).

74. J. P. Concordet, M. Haeussler, CRISPOR: intuitive guide selection for CRISPR/Cas9 genome editing experiments and screens. Nucleic Acids Res 46, W242–W245 (2018).

75. M. Bulajic et al., Differential abilities to engage inaccessible chromatin diversify vertebrate HOX binding patterns. Development, (2020).

76. A. M. Bolger, M. Lohse, B. Usadel, Trimmomatic: a flexible trimmer for Illumina sequence data. Bioinformatics 30, 2114–2120 (2014).

77. H. Li, R. Durbin, Fast and accurate short read alignment with Burrows-Wheeler transform. Bioinformatics 25, 1754–1760 (2009).

78. G. G. Faust, I. M. Hall, SAMBLASTER: fast duplicate marking and structural variant read extraction. Bioinformatics 30, 2503–2505 (2014).

79. S. Neph et al., BEDOPS: high-performance genomic feature operations. Bioinformatics 28, 1919–1920 (2012).

80. B. Langmead, S. L. Salzberg, Fast gapped-read alignment with Bowtie 2. Nat Methods 9, 357–359 (2012).

81. F. Ramirez, F. Dundar, S. Diehl, B. A. Gruning, T. Manke, deepTools: a flexible platform for exploring deep-sequencing data. Nucleic Acids Res 42, W187–191 (2014).

82. H. Li et al., The Sequence Alignment/Map format and SAMtools. Bioinformatics 25, 2078–2079 (2009).

83. J. T. Robinson et al., Integrative genomics viewer. Nat Biotechnol 29, 24–26 (2011).

84. A. R. Quinlan, I. M. Hall, BEDTools: a flexible suite of utilities for comparing genomic features. Bioinformatics 26, 841–842 (2010).

85. E. Riu, Z. Y. Chen, H. Xu, C. Y. He, M. A. Kay, Histone modifications are associated with the persistence or silencing of vector-mediated transgene expression in vivo. Mol Ther 15, 1348–1355 (2007).

86. D. Kim, B. Langmead, S. L. Salzberg, HISAT: a fast spliced aligner with low memory requirements. Nat Methods 12, 357–360 (2015).

87. D. Kim, J. M. Paggi, C. Park, C. Bennett, S. L. Salzberg, Graph-based genome alignment and genotyping with HISAT2 and HISAT-genotype. Nat Biotechnol 37, 907–915 (2019).

88. Y. Liao, G. K. Smyth, W. Shi, The R package Rsubread is easier, faster, cheaper and better for alignment and quantification of RNA sequencing reads. Nucleic Acids Res 47, e47 (2019).

89. M. I. Love, W. Huber, S. Anders, Moderated estimation of fold change and dispersion for RNA-seq data with DESeq2. Genome Biol 15, 550 (2014).

90. H. Wickham, in Use R*!*,. (Springer International Publishing : Imprint: Springer,, Cham, 2016), pp. 1 online resource (XVI, 260 pages 232 illustrations, 140 illustrations in color.

91. Y. Zheng, F. Ay, S. Keles, Generative modeling of multi-mapping reads with mHi-C advances analysis of Hi-C studies. eLife 8, (2019).

92. C. Levy-Leduc, M. Delattre, T. Mary-Huard, S. Robin, Two-dimensional segmentation for analyzing Hi-C data. Bioinformatics 30, i386–392 (2014).

